# Age and Gender-Dependent Group-Average Brain Biomechanics Models for Traumatic Brain Injury

**DOI:** 10.64898/2026.05.27.727945

**Authors:** Jiahao Wei, Ahmed Alshareef, Curtis Johnson, K.T. Ramesh

## Abstract

Experimental studies involving mechanical loading of the human head and brain in vivo are necessarily limited, making computational modeling essential for advancing our understanding of brain biomechanics. Demographic factors such as age and gender are known to influence brain anatomical structures, material properties, and potentially vulnerability to injurious loading. To address this, we construct six group-average brain models stratified by age and gender from a total of 135 subjects, and investigate the mechanical responses of these “group-average brains” using computational simulations. We use Pearson correlations to assess to what degree the group-average models represent individuals within each demographic category, showing strong correlations. Further, our p-value hypothesis test of the first principal strain across the six groups shows significant differences. This study demonstrates that age- and gender-stratified group-average models can effectively represent biomechanical responses of the individuals within the groups, and can reveal meaningful demographic differences that may influence susceptibility to traumatic brain injury. We show that the age-dependent change in material properties plays a greater role than anatomical changes in driving differences in the deformations. We hope to see increased utilization of these group-average models in both research and clinical applications.

## 1 Introduction

Traumatic brain injury (TBI) is a major cause of death and disability worldwide. At least 2 million cases of TBI are reported annually in the US alone, with varying degrees of severity [1] and significant societal and economic impact [2, 3]. Many experimental efforts have been devoted to studying the underlying pathophysiological mechanisms of TBI, generally relying on animal models such as primates [4] and rodents [5]. Ethical and safety concerns strictly limit human participation in such experiments, making computational modeling an essential methodology for studying TBI.

Magnetic resonance imaging (MRI) is a powerful imaging technique widely used in clinical practice for detecting conditions such as ischemic strokes, aneurysms, and tumors [6]. However, it is not ideal for the direct detection of mild traumatic brain injuries (TBIs) such as post-concussion syndrome and diffuse axonal injury [6]. MRI can also provide researchers with other valuable information about brain trauma, particularly in anatomical segmentation, and measure of motion and deformation. For instance, high-resolution segmentation of brain structures can be achieved by using automated algorithms applied to structural MRI [7, 8]. Brain motion can be analyzed using approaches such as tagged MRI (tMRI), DENSE MRI [9] and amplified MRI (aMRI) [10, 11], and (e.g.) harmonic phase MRI and tagged MRI (tMRI) enable reliable estimation of brain strain fields [12, 13, 14]. Diffusion-weighted imaging (DWI) provides insights into white matter tract orientations [15]. Additionally, brain biomechanical properties can be determined through magnetic resonance elastography (MRE) [16]. Overall, MRI not only informs subject-specific numerical models but also provides critical validation for such models.

Subject-specific full-field head and brain numerical models have been developed in recent decades. Kleiven analyzed 58 NFL cases using detailed finite element (FE) model of the human head [17]. Ji et al. demonstrated the significance of incorporating white matter anisotropy into FE models [18]. Zimmerman et al. built FE models and studied biomechanical signature of loss of consciousness [19]. While FE models are powerful, as pointed by Zhao and Ji, they suffer issues such as mesh locking and “hourglassing” [20], which encouraged researchers to build subject-specific material point method (MPM) models. Ganpule et al. developed subject-specific full-field MPM models and validated the models by comparing with deformation obtained from tagged MRI (tMRI) [21].

Incorporating the falx and tentorium, Lu et al. updated the MPM model to enhance the biofidelity [22]. Alshareef et al. integrated brain biomaterial properties from magnetic resonance elastography (MRE) into subject-specific, linear-viscoelastic (LVE) computational head models [23]. Upadhyay et al. upgraded the biomaterial constitutive relation to nonlinear-hyper-viscoelastic (NHVE) and compared with tMRI experimental observations [24] available in the Brain Biomechanics Imaging Resource (BBIR) database, available on the NeuroImaging Tools and Resources Collaboratory (NITRC) dataset [25]. In-vivo brain deformation during head motion has been investigated [26, 27, 28], and the high-fidelity data has been employed to validate the subject specific numerical models.

For both healthy and disease trajectories, there is value in developing group-average models that capture the biomechanical and physiological characteristics of distinct subpopulations. In brain studies, researchers have constructed imaging atlases for the human group-average cortex [29], rat group-average diffusion tensor imaging (DTI) [30], and group-level brain connectivity models [31]. In the context of traumatic brain injury (TBI), age and gender play critical roles in understanding the associated pathophysiological mechanisms [32], outcomes [33], prognosis [34], and mortality [35]. Although group-wise approaches in TBI research have been developing for years [18, 36], there is limited work on age and gender based group-average computational models. The concept of representativeness [37] is central to this approach, and thus requires a sufficiently large dataset which was not available until recently. Such models must provide reasonable and quantifiable accuracy with respect to the characteristics of the specific subjects they represent. Once we have established group-average computational models for TBI, we can interrogate the models in terms of commonly used metrics such as principal strains and axonal strains, and explore their potential clinical applications.

This work develops and assesses group-average computational models for traumatic brain injury, stratified across age and gender. We seek to address the following questions: (a) the degree to which the group-average model is representative of the individuals of the group (representativeness), (b) the distinctive biomechanical characteristics of each group, and (c) the use of the group-average models to consider potentially injurious loading. We expect that such insights will be useful both in terms of clinical applications as well as for the design of TBI mitigation approaches.

## 2 Methodology

Our overall approach is summarized in Fig. 1, which illustrates the overall workflow for this study. We use one group (mid-male, mid-age) as an exemplar. We first build the group-average brain computational model by defining group-average anatomy (a), developing the corresponding calibrated material properties (b), and applying the boundary condition (c) through the material points method (d). We then analyze the model outputs, such as the strain fields shown in (e). We validate the group-average models by comparing simulations results with experimental subject-specific tMRI strain from the BBIR dataset [25] (f). We then use Pearson correlation to demonstrate that the group-average models are representative of the individuals within the groups. We also compare the strains of different group-average models across age and gender under the same boundary condition at 45 ms to discuss their similarities and differences. Finally, we apply a hypothetical potentially injurious loading to the six group-average models to illustrate one possible use of these models.

**Figure 1:**
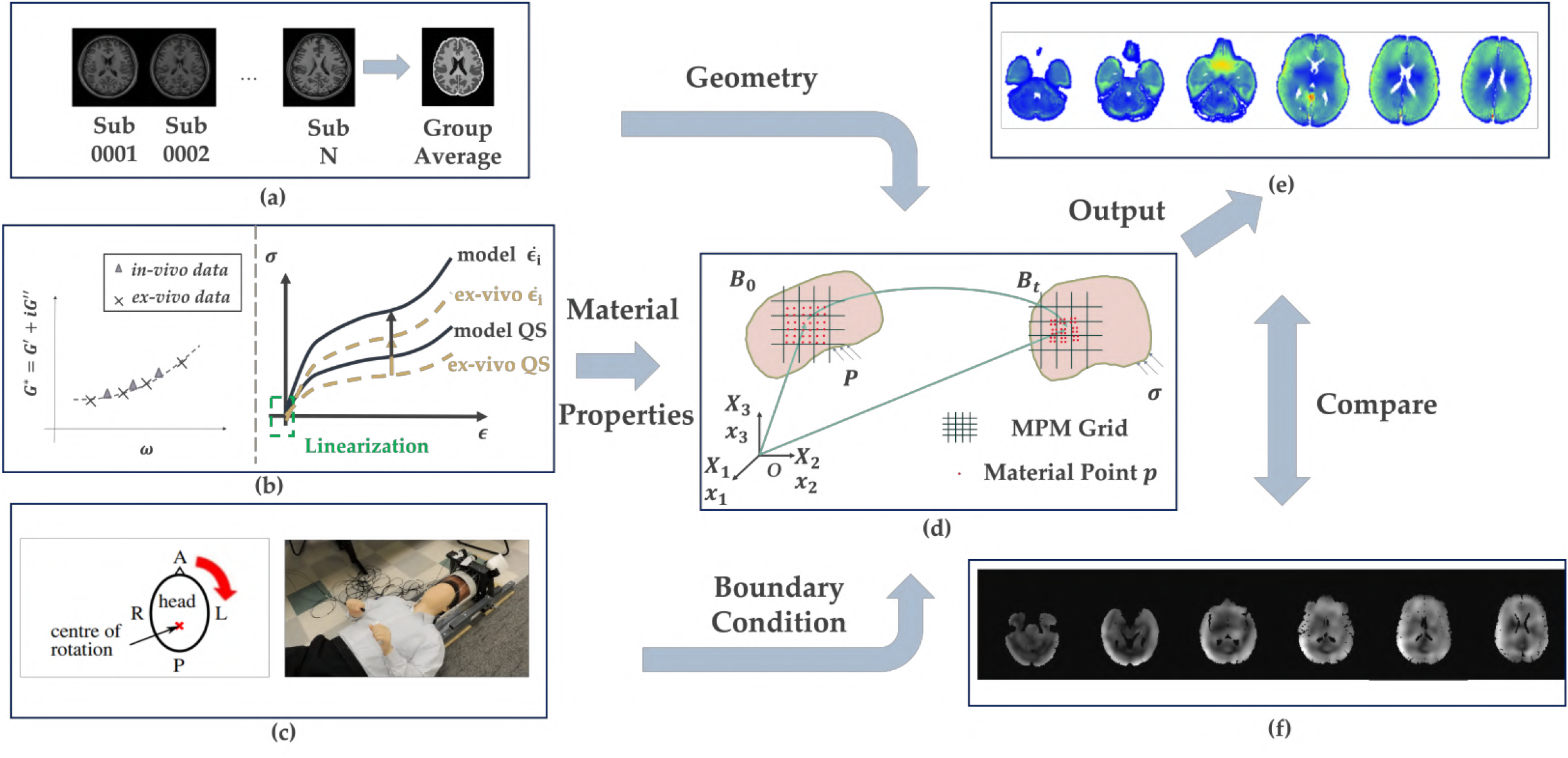
Our workflow for generation and assessment of group-average models. (a) N subjects of the group are registered in the standard space to obtain the group average anatomy. The obtained group average anatomy provides the geometry for the group average model. (b) In-ex vivo material properties calibration. Calibrated material properties inform material parameters of different brain segments of the group average model. (c) A picture taken from a neck rotation experiment with conceptual plot. Volunteers’ necks rotate about superior-inferior axis, as denoted in the red cross mark. Collected rotational angular velocity offers kinematic boundary condition of the group average model. (d) Material point method (MPM). Computational information carried by discrete Lagrangian material points is mapped to Eulerian grids, updated on the grids, and mapped back to Lagrangian material points within each time step. (e) A strain spatial distribution of the group average MPM model on six transverse slices. (f) A strain spatial distribution generated by harmonic finite element method (HARP-FE) from tagged MRI (tMRI) data of subject 0001 (mid male, age=31 from BBIR database).

### 2.1 Defining group-average anatomy

A total of 135 high-resolution subject specific anatomies were obtained from volunteers through structural MRI scans such as T_1_, T_2_ weighted, diffusion weighted imaging (DWI), and susceptibility weighted imaging (SWI). Details about acquisition and parameter settings can be found in [26], [38], and [39]. These scans enable sufficient contrast among soft brain components to allow segmentation through automatic algorithms [40]. Each of the models comprises the following 12 segments at a 1.5 mm voxel resolution: brainstem, cerebellum gray, cerebellum white, cerebrum white, cortical gray, cerebrospinal fluid (CSF), deep gray, falx, foramen, subarachnoid space (SAS), skull, and ventricles. The segmented 132 spatially localized atlas network tiles (SLANT) [38] are converted to 8-label anatomical structures, and exported as point files for our material point simulations [41].

As shown in Table 1, we first divide the subjects into six groups based on their age and gender, and denote each group as follows: young female, young male, mid female, mid male, older female, and older male. The number of available magnetic resonance imaging (MRI), magnetic resonance elastography (MRE), and tagged magnetic resonance imaging (tMRI) datasets in each group are also shown in the table. The MRI data is used for obtaining the group-average anatomical structure, which is briefly described in this section. The MRE data is used to calibrate group-average material properties, which is introduced in section 2.3. The tMRI data, also called experimental data in the following text, serves as the ground truth for comparing the group average computational model with the experiment results. Note that we have 0 tMRI scans for the young female group. However, for the purposes of this effort, 6 female subjects (0013, 0016, 0018, 0019, 0023, and 0035) whose ages are under 25, have been treated as part of this young female cohort (see also [38]). Further, we focus on a head rotation boundary condition, and so we do not use the tMRI data from some subjects that only had data for an extension type of kinematic boundary condition applied during the scan.

**Table 1:**
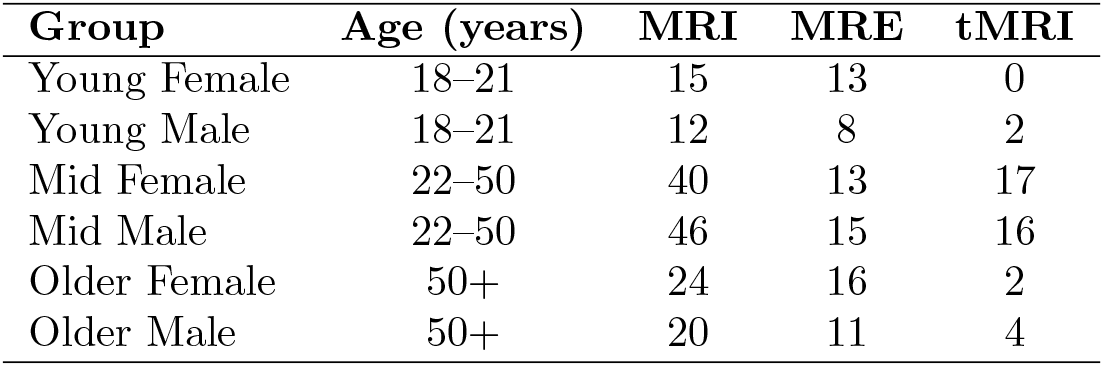
Grouping based on age and gender.

| Group | Age (years) | MRI | MRE | tMRI |
| --- | --- | --- | --- | --- |
| Young Female | 18-21 | 15 | 13 | 0 |
| Young Male | 18-21 | 12 | 8 | 2 |
| Mid Female | 22-50 | 40 | 13 | 17 |
| Mid Male | 22-50 | 46 | 15 | 16 |
| Older Female | 50+ | 24 | 16 | 2 |
| Older Male | 50+ | 20 | 11 | 4 |

To acquire the group average anatomy, all individual subject images are first rigidly registered to the standard MNI 152 space using the Advanced Normalization Tools (ANTs) [42]. Given the subject-specific anatomical information, group-average anatomical structures are then generated (Figure 1 (a)) by nonlinearly registering all subjects within each group using the ANTs template creation script with multimodal inputs. This is used to create average T_1_ weighed and T_2_ weighted images, and brain segments for each group, in this sense forming a group-average anatomical structure. Similar to the subject-specific brain segmentation, group average brain segmentations are acquired through SLANT and combined into 8 brain segments. In addition, we manually segmented the cerebrum white matter into corona radiata and corpus callosum using ITK-SNAP [43]. Further details about the generation of the group-average anatomical structures are provided elsewhere [38].

Note that fiber directions and fractional anisotropy (FA), which are included in the subject-specific brain simulations [24], are not available in the group-average models due to limitations in the image registration algorithms. While our current registration algorithms do not fully preserve subject-specific white matter fiber directions and FA, we expect that future advancements may enable group-average simulations to include fiber orientations and anisotropy.

### 2.2 Determining group-average material properties

Our computational model assumes that the material behavior, defined at each material point, is based on either linear viscoelasticity or nonlinear hyper-viscoelasticity [24]. Two major steps are involved in the calibration (Figure 1(b)) process: first, calibration of the linear viscoelastic material properties (LVE), and second, calibration of the nonlinear hyper-viscoelastic (NHVE) material properties.

Details about subject-specific [23] and group-average linear viscoelastic material properties are presented elsewhere [38], and summarized as the following. In the LVE calibration, the material properties are defined in terms of Prony series. LVE calibration uses both *in-vivo* and *ex-vivo* data [23]. The *in-vivo* experimental data is acquired from MRE experiments at frequencies of 30Hz, 50Hz, and 70Hz using a resonant pneumatic actuator system with a soft pillow driver that induces displacements [27]. In the time domain, the shear modulus *G* (*t*) is:

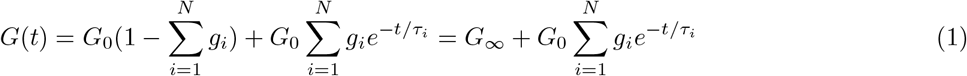

Here,*G*_0_ is instantaneous (short-term) shear modulus, and *µ*_*∞*_ is long-term shear modulus. *g*_*i*_ and *τ*_*i*_ are fractional shear contributions and fractional time constants of the i-th Prony Series in the time domain. However, in the MRE scan, the moduli are obtained in the frequency domain as storage modulus *G*′ and loss modulus *G*″ at 30, 50, and 70 Hz [27]. The moduli in the time domain and frequency domain are related by the following relationships:

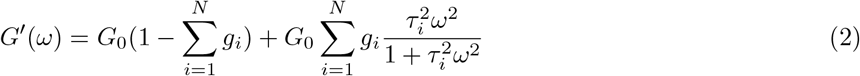

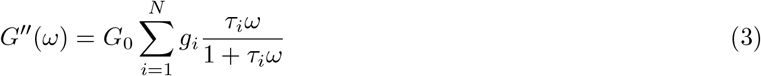

To obtain group-average moduli and fractional time constants in the time domain, subject-specific MRE images are first nonlinearly registered to the group-average T2w image. The associated transformations are used to warp the subject-specific MRE moduli to the group-average space at the three MRE harmonic loading frequencies. Group average MRE moduli and damping ratio in the frequency domain are computed by averaging voxel-wise values of all the specific subjects of each group with associated standard deviation. Using the MRE moduli and damping ratio at the three actuation frequencies, and ex-vivo data at other frequency values [44], an optimization scheme is implemented that scales ex-vivo storage/loss modulus versus frequency curves to in-vivo MRE data points using a fitted scaling factor. In the scheme, the scaling factor is acquired by extrapolating the limited MRE frequencies (30, 50, 70 Hz) to a wider range of frequencies using root mean square errors. The root mean square error across the entire frequency range is used in the optimization scheme to fit the Prony series parameters [38].

The LVE group-average models are simulated with these fitted parameters, while a second calibration step uses the fitted Prony series parameters to further calibrate the properties of the visco-hyperelastic O-USS constitutive relation. Details about the motivation, implementation, and significance of the O-USS model are provided in [45, 24]. The O-USS constitutive relation describes the visco-hyper-elastic attributes of the brain tissues. In the O-USS constitutive relation, the Cauchy stress tensor is related to the deformation tensors following equation (4).

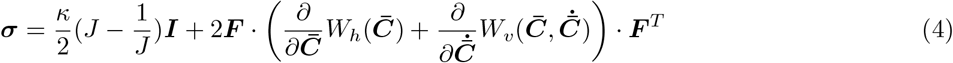

Here,***F*** is the deformation gradient tensor, 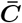 is the modified right Cauchy-Green tensor, where 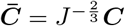 and *J* = det(***F***). *W*_*h*_ denotes the Ogden energy density function, expressed as:

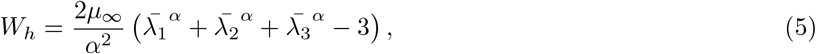

and *W*_*v*_ is the USS dissipative strain energy density function:

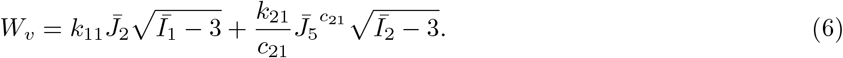

Here *µ*_*∞*_ represents the long-term shear modulus, *α*represents the compression-tension asymmetry, and 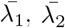 and 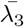 are the principal stretch ratios of 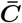. *W*_*v*_ is the USS dissipative strain energy density function, where *k*_11_, *k*_21_, and *c*_21_ are linear sensitivity control parameter, nonlinear sensitivity control parameter, and rate sensitivity index, respectively. Note that to satisfy continuum thermodynamics, we need to constrain these parameters so that *k*_11_ >0, *k*_21_ >0, and *c*_21_ >0.5. The invariants in equation 6 are: 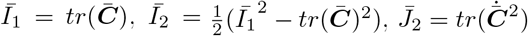, and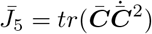.

Nonlinear visco-hyperelastic (NVHE) calibration also uses both *in-vivo* and *ex-vivo* data. The calibrated *G*_0_ */G*_*∞*_, *g*_*i*_, and *τ*_*i*_ from the LVE calibration step are plugged in equation (7) to obtain strain-rates dependent apparent moduli *G*_*app*_ and *E*_*app*_, using the strain rates 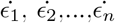, under small strain regime (*ϵ*_*LV E*_ <0.01).

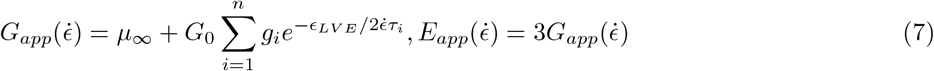

Under the same small strain regime, we assume that the dependence of nonlinearity of the material moduli is negligible. Thus, the corresponding moduli represented in the O-USS model, called tangent moduli *G*_*tan*_ and *E*_*tan*_, should have the same values as the LVE model. The tangent moduli *G*_*tan*_ and *E*_*tan*_, corresponding to the “linearization” regime in Fig. 1 (b), are expressed in the O-USS model as the following [24]:

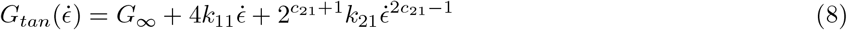

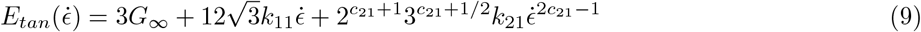

Equating the tangent moduli with the apparent moduli, we can calibrate the parameters *k*_11_, *k*_12_ and *c*_21_ in the linear regime, denoted as 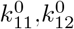 and 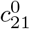. These parameters provide initial values for moduli under finite strain regime. Note that *k*_11_ >0, *k*_21_ >0, and *c*_21_ >0.5 as set forth.

Compression-tension asymmetry parameter *α*in equation (5) is calibrated using literature large-deformation data points [24]. At this point, we have successfully known the following parameters: 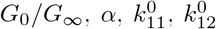 and 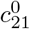, among which 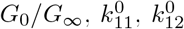 and 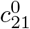 are purely informed by small-strain data. In order to preserve the biofidelity and calibrate the O-USS model under large deformation regime, an optimization problem is designed.

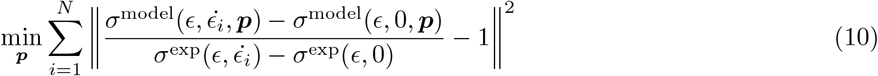

The target function of the optimization in expression (10) contains the L2 norm of the difference between the overstress of the O-USS model and the overstress of ex-vivo data. Here, overstress is defined as stress at certain strain rates 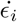subtracted by stress at quasi-static state. The goal of the optimization is to minimize L2 norm toward to zero. The initial guess of the vector***p***= [*k*_11_, *k*_12_, *c*_21_] is taken 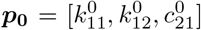, with ±10% inequality constraint on tangent module *G*_*tan*_ in equation (8). The optimization step will give values of 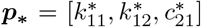. In a later step, voxel-wise 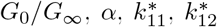 and 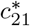 from the same brain regions will be averaged within each segment. The bulk modulus *κ* is obtained from the literature [46], and the value enforces incompressibility. Together, they will be used as material parameters to build the MPM simulation model. Table 2 shows the mid-male group average model parameters used for its MPM model. The O-USS material parameters for other groups can be found in the Appendix A. The same material parameters are used across all the group average MPM models for the sub-arachnoid space, falx and tentorium, skull, ventricles, and CSF.

**Table 2:** Mid-male group average material parameters.

| O-USS material parameters |  |  |  |  |  |  |  |
| --- | --- | --- | --- | --- | --- | --- | --- |
| Brain Substructure | $G_\infty$ (Pa) | $\alpha$ | $k_{11}$ (Pa·s) | $k_{21}$ (Pa·s<sup>2-2c<sub>21</sub></sup>) | $c_{21}$ | $\kappa$ (GPa) | $\rho$ (kg/m<sup>3</sup>) |
| Brain Stem | 938.1 | 2.61 | 2.0 | 357.6 | 0.571 | 1.46 | 1040 |
| Cerebellar Gray | 732.2 | 2.71 | 10.7 | 324.7 | 0.636 |  |  |
| Cerebellar White | 1026.4 | 2.71 | 9.1 | 473.8 | 0.611 |  |  |
| Corona Radiata | 1209.2 | -3.47 | 5.5 | 624.9 | 0.610 |  |  |
| Corpus Callosum | 1541.6 | -2.32 | 4.5 | 576.2 | 0.565 |  |  |
| Cortical Gray | 967.4 | -3.76 | 8.8 | 453.4 | 0.631 |  |  |
| Deep Gray | 1551.4 | 4.92 | 6.8 | 667.3 | 0.584 |  |  |
| Linear viscoelastic material parameters |  |  |  |  |  |  |  |
| | $G_0$ (kPa) | $g_1$ | $\tau_1$ (ms) | $\kappa$ (GPa) | $\rho$ (kg/m<sup>3</sup>) | Reference | |
| Sub-arachnoid space | 0.5 | 0.8 | 12.5 | 1.46 | 1133 | [46, 24, 47] |  |
| Linear Elastic material parameters |  |  |  |  |  |  |  |
| | $E$ (MPa) | $\nu$ | $\rho$ (kg/m<sup>3</sup>) | Reference | | | |
| Falx and Tentorium | 31.5 | 0.45 | 1133 | [48, 24] |  |  |  |
| Skull | 8000 | 0.22 | 2070 | [49, 24] |  |  |  |
| Newtonian Fluid |  |  |  |  |  |  |  |
| | $\mu$ (Pa·s) | $\gamma$ (N/m<sup>3</sup>) | $\kappa$ (GPa) | Reference | | | |
| Ventricles and CSF | 1.0e-3 | 7.15 | 1.46 | [50] |  |  |  |

### 5.3 Simulations using the material point method

The Material Point Method (MPM) is a particle-in-cell computational method that combines both Lagrangian and Eulerian frameworks to solve partial differential equations [51, 52]. In the Lagrangian framework, MPM discretizes a continuum into material points, with each point carrying essential physical information such as mass, volume, position, velocity, deformation gradient tensor, external stress, and internal stress. At time *t+*_*n*_, the essential information carried by all material points are assumed to be known. The known information is then mapped onto a uniform Eulerian grid using a mapping matrix **S** and its gradient ∇**S**. On the nodes of the Eulerian grid, acceleration is computed by solving the discrete momentum equation derived from its weak form. Once the acceleration is determined, the updated nodal velocity vectors at time *t*_*n*+1_ are obtained. This updated nodal velocity vectors are then mapped back to the material points in the Lagrangian framework using the gradient of the shape function ∇**S**, allowing the computation of the velocity gradient ∇**v** in the Lagrangian space. The updated velocity gradient is used to compute the Lagrangian deformation gradient tensor at time *t*_*n*+1_ through a multiplicative incremental deformation gradient tensor. Similarly, other quantities such as volume, position, velocity, and stresses (both external and internal) are updated following an analogous procedure [53]. For whole-scale brain simulation, MPM offers several advantages, including the ability to simulate multi-phase systems, ease of converting MRI anatomical structures into MPM-compatible geometries, and the prevention of mesh locking [21, 24].

We use MPM to compute group average brain deformations under rotational kinematic boundary conditions of the whole head using the geometries (group average anatomical structures) defined in Section 2.1, and using the material properties defined in Section 2.2. Our full-scale group-average MPM brain simulations use a grid resolution of 96 *×*96 *×*72 in a 19 cm by 10.6 cm by 16.4 cm 3D box space. The temporal duration is 189 ms, with a Courant–Friedrichs–Lewy (CFL) factor of 0.4. The simulations are multi-phased: for instance, ventricles and CSF are treated as fluids, while skull and cerebrum outer gray are treated as linear elastic and visco-hyper-elastic solids. For this work, all jobs ran for approximately 55 hours on the Johns Hopkins University high performance computing cluster Rockfish using 720 CPUs (15 nodes, each with 48 CPUs).

### 2.4 Initial and boundary conditions

Initial and boundary conditions are required to run simulations using the computational model. In this study, kinematic boundary conditions for neck rotation about the inferior–superior (IS) axis are obtained during the experiments from an angular position sensor integrated into a custom MRI-compatible rotation device. For illustration, Figure 1(c) shows a volunteer (not one of our subjects) using the custom rotation device inside a 3T Siemens mMR Biograph MRI scanner. This device is designed to restrict rotational motion to sub-injurious levels (2–3 *g*or 150–350 *rad/s* ^2^) [27], and the recorded rotational motion of the head is used directly for our simulations. The specific boundary conditions applied here (as an example) come from volunteer subject 0001 (male, age=31) scanned at NIH under a protocol NIH approves. During data acquisition for subject 0001, the device was operated manually, with the head initially leaned toward the right shoulder. Head rotation is initiated by releasing a latch, resulting in an initial angular acceleration followed by mild deceleration determined by the device’s mechanical constraints. This motion is centered on the head’s IS axis, as depicted schematically in the inset of Figure 1(c).

### 2.5 Comparing Simulations with Experimental Data

In the experiments, subject-specific 3D strain fields are obtained through multi-slice tagged MRI (tMRI) during repetitive loadings using the same MRI-compatible device. tMRI, combined with harmonic finite element method (HARP-FE), which ensures geometrical integrity and compatibility, provides accurate 3D strain measurements [54]. All of the associated data is available in the BBIR dataset [25]. We consider this experimental data to represent ground truth for determining the representativeness of our group average models.

We begin by comparing the computed strain histories from our group-average models with the individual experimental strain histories that have been post-processed by HARP-FE [54] for subjects within the corresponding group. There are two aspects that we address in these comparisons: the *peak time* (time at which the first principal strain E_1_ of Lagrangian Green strain tensor **E** achieves its maximum value), and the evolution of the principal strains with time (*principal strain histories*, e.g. the magnitude of E_1_ over time), from which we can tell whether the group average model responses evolve similarly with time asthe responses of the individuals in the group.

The comparisons are conducted in terms of selected brain segments. Figure 3 presents six transverse planes within the MPM simulation space that we use to demonstrate the brain anatomy (this set is for the mid-male group). These six transverse planes have IS coordinates of −35 *mm*, − 30 *mm*, −27 *mm*, 0 *mm*, 10 *mm*, and 15 *mm*. We will focus on the two transverse planes with the IS coordinate of 15 mm and −35 mm of mid-male for much of the analysis. The 15 mm transverse plane (denoted as the *cerebrum slice* from now on) is chosen because it encompasses all key brain cerebrum components of interest, namely cerebrum white matter (including corona radiata and corpus callosum), cerebrum outer matter, and deep gray matter. The −35 mm transverse plane (denoted as the *cerebellum/brainstem slice*) includes cerebellum white matter, cerebellum gray matter, and brainstem. Together, the six transverse planes provide an anatomical representation of the brain (in this case, the mid-male brain). As for metrics of comparison, we choose the 50th and 95th percentile of the first principal strain E_1_ of the Lagrange-Green strain tensor across all voxels within each brain segment. Such segment-wise percentile values are called *segmental strain histories* in the text, and are collected every 3 *ms* from the simulations.

We use the Pearson correlation *r* (equation 11) between the segmental strain histories and the experimental individual (subject-specific) segmental strain histories to provide a quantitative metric of the degree to which the GAM represents the members of the group:

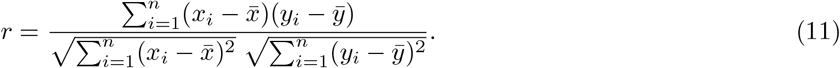

Note that our group-average geometries are registered and transformed to the common MNI 152 template space [38], while the group’s individual geometries are obtained in their own spaces. Thus, the corresponding cerebrum slice and cerebellum/brainstem slice of the individual experimental data do not share the identical IS coordinates of −15 *mm* and 35 *mm*. To determine their corresponding IS coordinates in the subjects, we manually extracted the two corresponding transverse planes that share the most similar IS locations, anatomical shapes, and segmental sizes to the best extent, using ITK-Snap [43]. When comparing our group average model with the subjects of the group, it is also important to remember that the kinematic boundary conditions used for the group-average simulation is obtained from the case of subject 0001, and differs from the actual kinematic boundary conditions for other specific subjects (although the loading platform enforces consistency of direction, not the magnitude).

The common features of spatial strain distribution are also analyzed on the six transverse planes and the corresponding transverse planes of subject 0001 (see above for how we select these planes) at 27 *ms*, 45 *ms*, and 63 *ms*. The selected time points include times before and after the maximum principal strains are achieved at approximately 45 ms. The 18 *ms* time difference is chosen to match the time gap between each experimental data collection. Our analysis here focuses on the spatial distribution of the first principal strain E_1_ as an exemplar, and we use this to compare the group-average models and the experimental data.

## 3 Results and Discussion: Assessing our Group-Average Models

We begin by noting that the ability of our computational approach to capture the brain biomechanics for individual subjects has been demonstrated in previous work [24, 21], and so validation of the approach itself has already been completed. Our focus in the next few subsections is (a) first on demonstrating that these Group-Average Models (GAMs) are representative of the behaviors of the members of the group, (b) next on comparing the biomechanical responses of the groups to clarify age and gender differences, and finally (c) assessing the consequences of applying a fictitious injury-causing loading to each of the groups. All the demonstrations, comparisons, and extended applications will be discussed in terms of the brain anatomical segments defined in Figure 3.

### 3.1 How representative of the group is a Group-Average Model?

We use the mid-male group as an exemplar for the purpose of discussing how well a given group-average model (GAM) represents the members of the group. The descriptive metrics we choose are based on the principal strains in these isotropic materials: first, the maximum principal strain E_1_ is commonly used as a metric to determine the threshold for injuriy [55, 56]; and second, the maximum shear strain (another commonly used metric for the injury threshold [57]) can be computed easily from the three principal strains.

The time-history of the first principal strain (E_1_), as computed from the full mid-male group average NVHE model, is presented in Figure 4 for three (cerebrum deep gray, cerebrum outer gray, and cerebrum while) of the segments described in Figure 3, showing the results for the *cerebrum slice*. Results are shown as solid blue lines for a timescale up to just over twice the time of which the rotation is applied (see Figure 2). Since the first principal strain is a field variable, on the cerebrum slice, we extract voxel-wise E_1_ values of each brain segment at each time, calculate the 50th and 95th percentile E_1_ values of each extracted brain segment, and plot these values as a function of time. Note that the group average data coming from the stored computational output starts at 0 ms and is picked every 3 ms. Both the 50-th (top row) and 95-th (bottom row) percentile results show that the first principal strain in each segment first increases with time over about 50 milliseconds, then decreases before beginning to rise again for a small second peak. Each of the three segments presented has a different strain history, with the computed strain during the first oscillation being lowest in the cerebrum deep gray segment and highest in the cerebrum outer gray segment.

**Figure 2:**
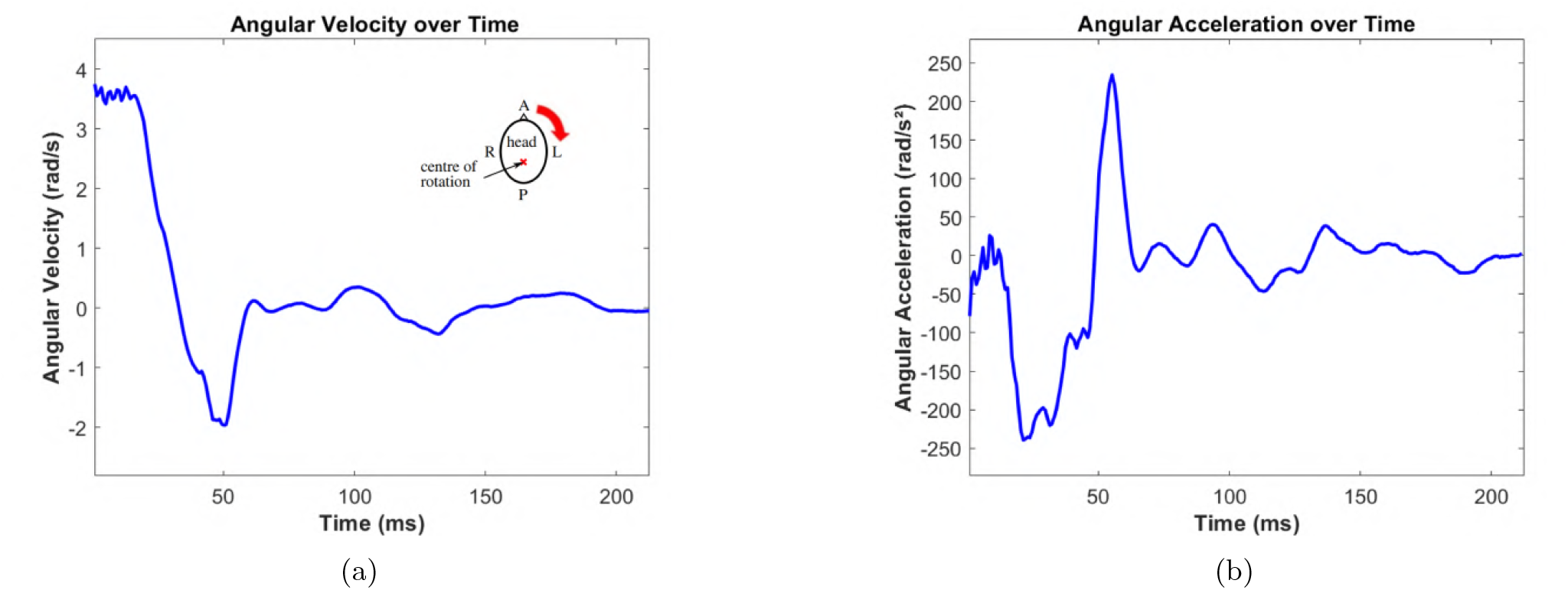
Example boundary conditions applied during our simulations. Angular velocity (a) and angular acceleration (b) for the head as functions of time as measured for subject 0001.

**Figure 3:**
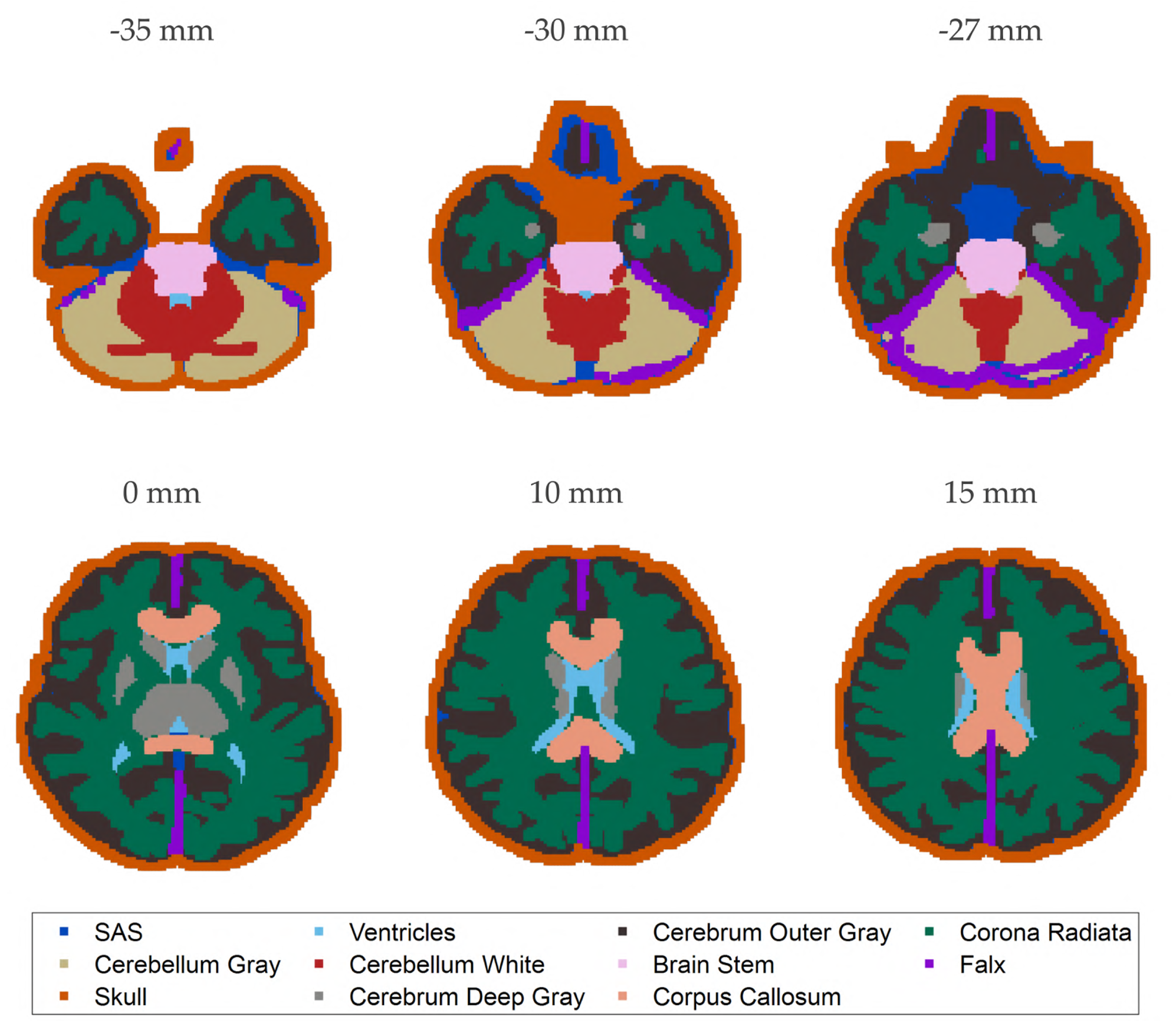
Six selected anatomical transverse planes of mid-male group with interior-superior (IS) coordinates of −35 mm, −30 mm, −27 mm, 0 mm, 10 mm, and 15mm.

**Figure 4:**
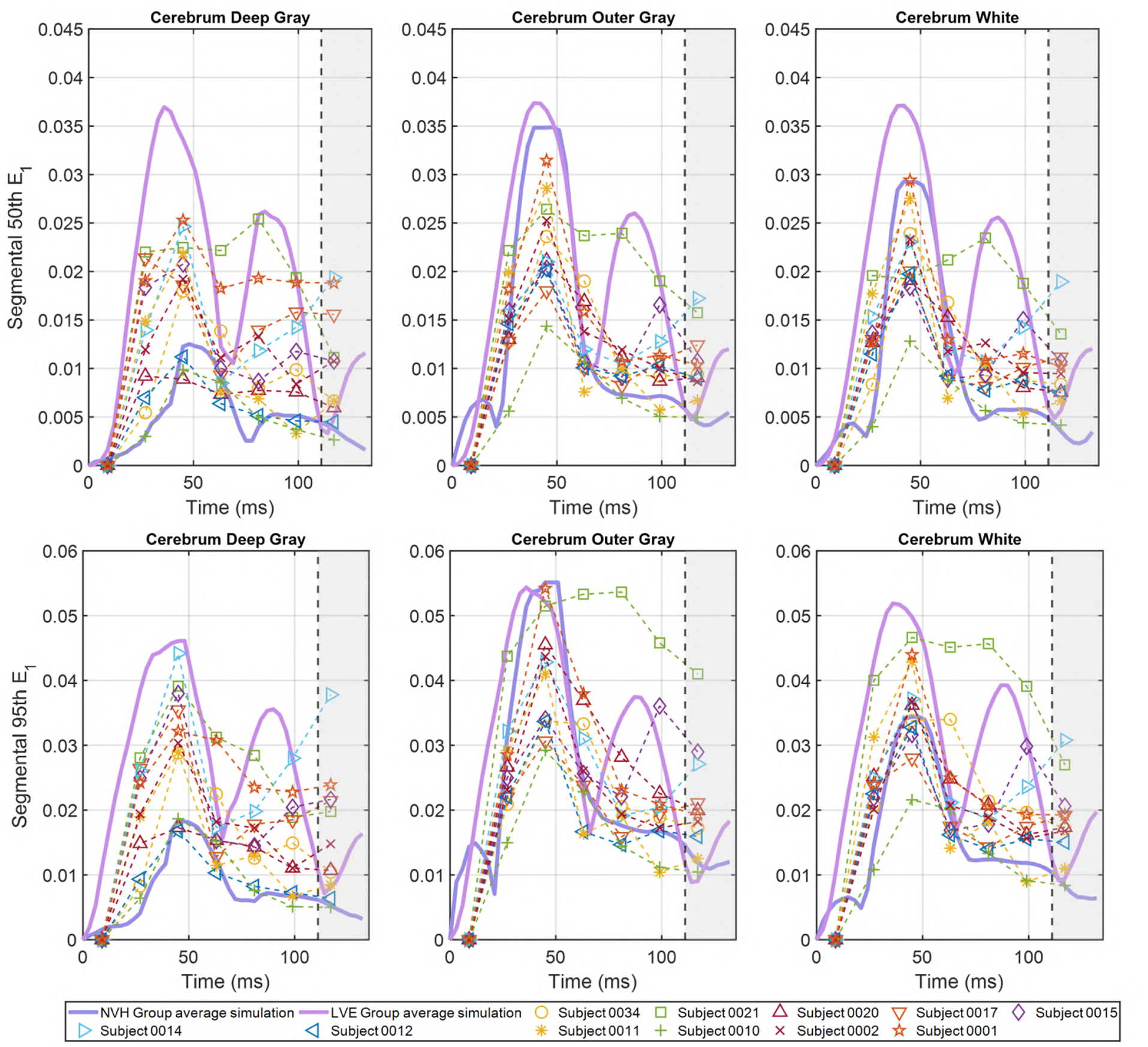
Segmental strain histories on IS = 15 mm (*cerebrum slice*). 50th (first column) and 95th (second column) percentile first principal strains E_1_ temporal variation of cerebrum segments - cerebrum deep gray, cerebrum outer gray, and cerebrum white - are presented. The solid blue lines are E_1_ of mid-male group average simulation model using nonlinear visco-hyperelastic model. The solid purple lines use linear visco-elastic model. The dashed lines are experimental data E_1_ obtained from subject-specific tMRI data. Note that at each time step, the 50th and 95th percentile E_1_ are not necessarily extracted from the same material point.

The corresponding individual (subject-specific) experimentally measured tMRI strain histories are also presented in Figure 4. There are 11 that start at 9 ms, collected approximately every 18 ms. The 9 ms offset is because the first frame of the tMRI image data collection is not completed until 9 ms [24]. The time-resolution of the experimental data is lower than than of the computational data. The experimental data points beyond 100 ms are shaded because they are considered less reliable due to tag fading [58]. Note the range of responses obtained within these 11 subjects. Each subject of course has his own specific anatomical structures, and subject-specific material properties. Further, each subject has his own kinematic boundary condition which is different but in general consistent (due to the rotational device setup) from that used for the mid-male group average model simulation. The particular boundary condition used in the simulation is that of subject 0001, and that subject’s experimental data is represented by the red stars in Figure 4.

The times at which the peak strain is achieved are about 45 *ms* for the NVHE group-average model for all of the segments in Figure 4, and this is also the time when the peak strain is observed in the experimental data for most subjects (recognizing the more discrete nature of the experimental data). The magnitude of the peak strain in the NVHE group-average simulations is different for the three segments shown, and this is also true of the experimental data. The overall computed time history of these strains falls within the general range of the histories of the experimental data (noting here that the time resolution is much lower for the experiments). A very similar comparison of the computed mid-male group-average model and the experimental data for these 11 subjects is observed for the *cerebellum slice* in Figure 5. Note that the experimentally determined strains for some subjects (e.g. subject 0017 in the cerebellum white segment) can be quite different from the rest, demonstrating the range across the subjects.

**Figure 5:**
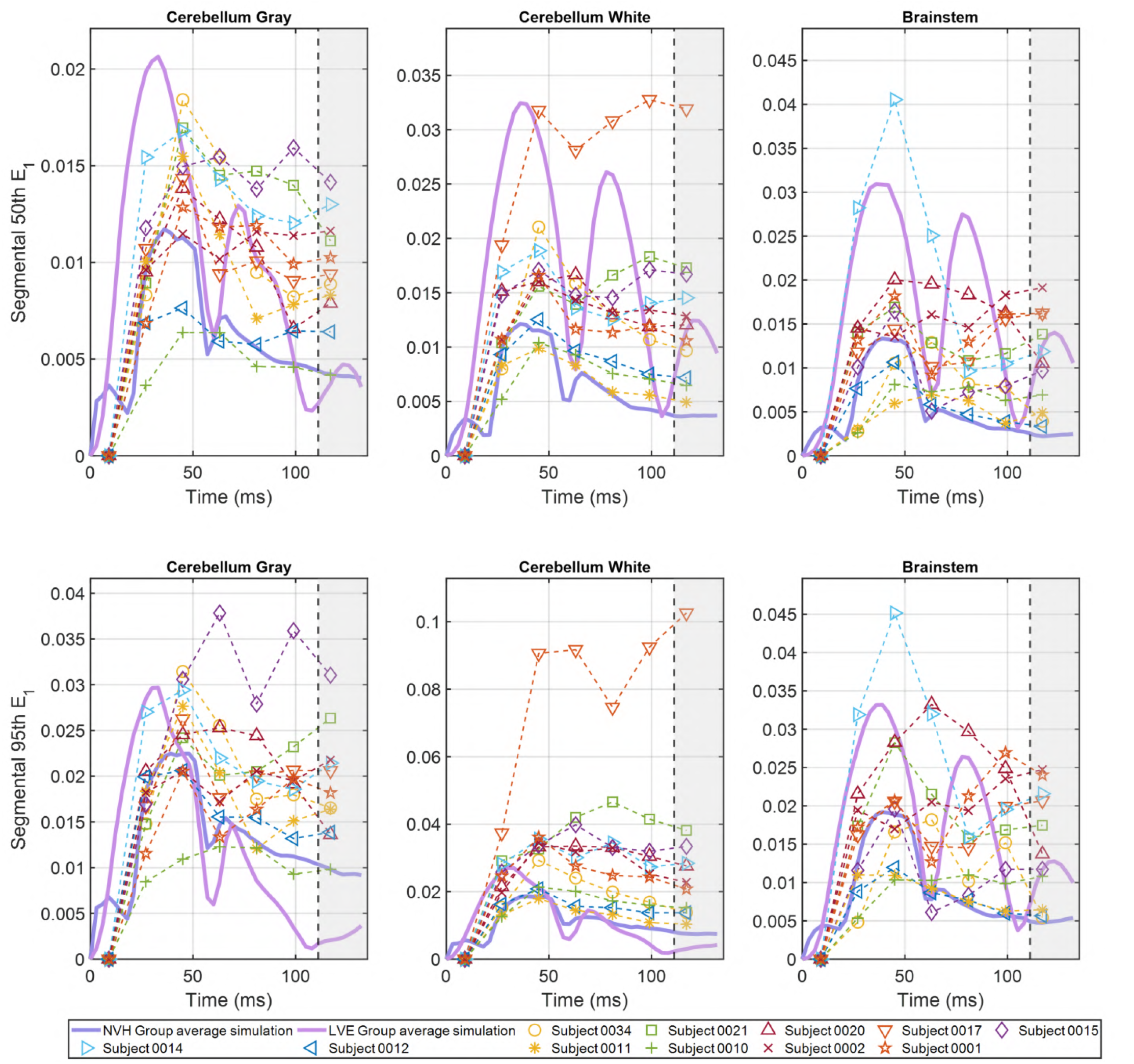
Segmental temporal variation comparison on IS = −35 mm (*cerebellum slice*). 50th (first column) and 95th (second column) percentile first principal strains E_1_ temporal variation of cerebellum and brainstem. The solid blue lines are E_1_ of mid-male group average simulation model using nonlinear visco-hyperelastic model. The solid purple lines use linear visco-elastic model. The dashed lines are experimental data E_1_ obtained from subject-specific tMRI data. Note that at each time step, the 50th and 95th percentile E_1_ are not necessarily extracted from the same material point.

In Figure 4, considering the cerebrum outer gray matter located near the skull, both the 50th and 95th percentile E_1_ group-average values resemble the experimental E_1_ values of subject 0001. This result is expected, as the principal strain E_1_ in the cerebrum outer gray matter is strongly influenced by the applied kinematic boundary conditions due to its proximity to the brain’s outer regions. That said, in other segments, we should expect some differences between the group average simulations and the response of subject 0001 for several reasons: different material properties; the fact that the HARP-FE algorithm [54], in the post-processing of tMRI data, incorporates hyperelastic behavior yet neglects time-dependent viscous internal dissipation [54]; and the differences between the GAM anatomy and subject-specific anatomy. Cumulative internal viscous dissipation may reduce strain magnitudes as energy is transferred from outer to inner brain regions, e.g. from cerebrum outer gray to cerebellum white matter.

Both Figure 4 and Figure 5 show that all experimental data curves exhibit at least one peak at approximately 45 ms, where the 50th or 95th percentile E_1_ values increase from 9 ms to 45 ms and then decrease. A similar trend is observed in the NVHE group-average simulation data. In subject 0001, subject 0015, and subject 0034, they present a second increase-decrease peak at approximately 100 ms in the cerebrum deep gray, outer gray, and white. However the experimental data percentile E_1_ values beyond 100 ms (shaded regions in Figure 4 and Figure 5) are less reliable due to limitations in imaging algorithms [58]. On the simulation side, the NVHE group-average curves show an early local peak at about 10 *ms* in all segments except the cerebrum deep gray. We do not currently understand the source of this early peak.

Overall, the NVHE mid male group-average results (blue lines) in Figure 4 and Figure 5 appear to agree reasonably well with the range of the corresponding individual tMRI experimental data in terms of peak time and strain magnitudes, but quantitative comparisons are of course difficult to make by eye. Thus, we make a direct comparison (using Pearson correlation) of the 95th percentile E_1_ from the mid-male group-average model with the experimental results of all the mid-male subjects in Figure 4 and Figure 5 across cerebrum and cerebellum brain segments. To compute the correlation, we take the first 7 time points from the experimental data before 100 ms, find the corresponding 95th percentile E_1_ values from the simulation at those times, and use a Matlab built-in correlation function for the computation.

Figure 6 shows strong [59] correlations (with five out of six segmental median correlations greater than 0.7) between the mid-male GAM and the mid-male experimental data. We also highlight the Pearson correlation with subject 0001 using the red circles in the figure, as it shares the same kinematic boundary condition as the group-average simulation. We see that in the cerebrum region, which makes up approximately 88% of the brain volume [60], strong correlations are presented, with correlation medians greater than 0.75 in all segments. Particularly, the segmental correlations between the mid-male group-average model and subject 0001 are strong (*r* >0.7) in cerebrum and cerebellum white, with cerebrum white very strong [59] (*r* >0.9). The reader is referred to Figure 12 in Appendix B for mid-female correlations: the correlations are strong in cerebrum outer gray, cerebrum white, and cerebellum white; and moderate (0.5< *r* <0.7) in cerebrum deep gray, and cerebellum gray. Note that there is no experimental subject in the mid-female group sharing the same boundary condition with the mid-female group average model. We also studied the correlations between the older-male group average model and the 4 experimental subjects in the group (see Table 1). The median correlations of cerebrum deep gray and cerebrum white are close to very strong (*r* >0.85), with cerebrum outer gray moderate (*r* = 0.579). Correlation analyses are omitted for the rest of the groups since their tMRI data is either scarce (with less than 1 rotational kinematic boundary condition) or contains cadaver only (in the case of the older female group). In the sense of Pearson correlations, our group average models are good representatives of the members of the group, at least in terms of E_1_.

**Figure 6:**
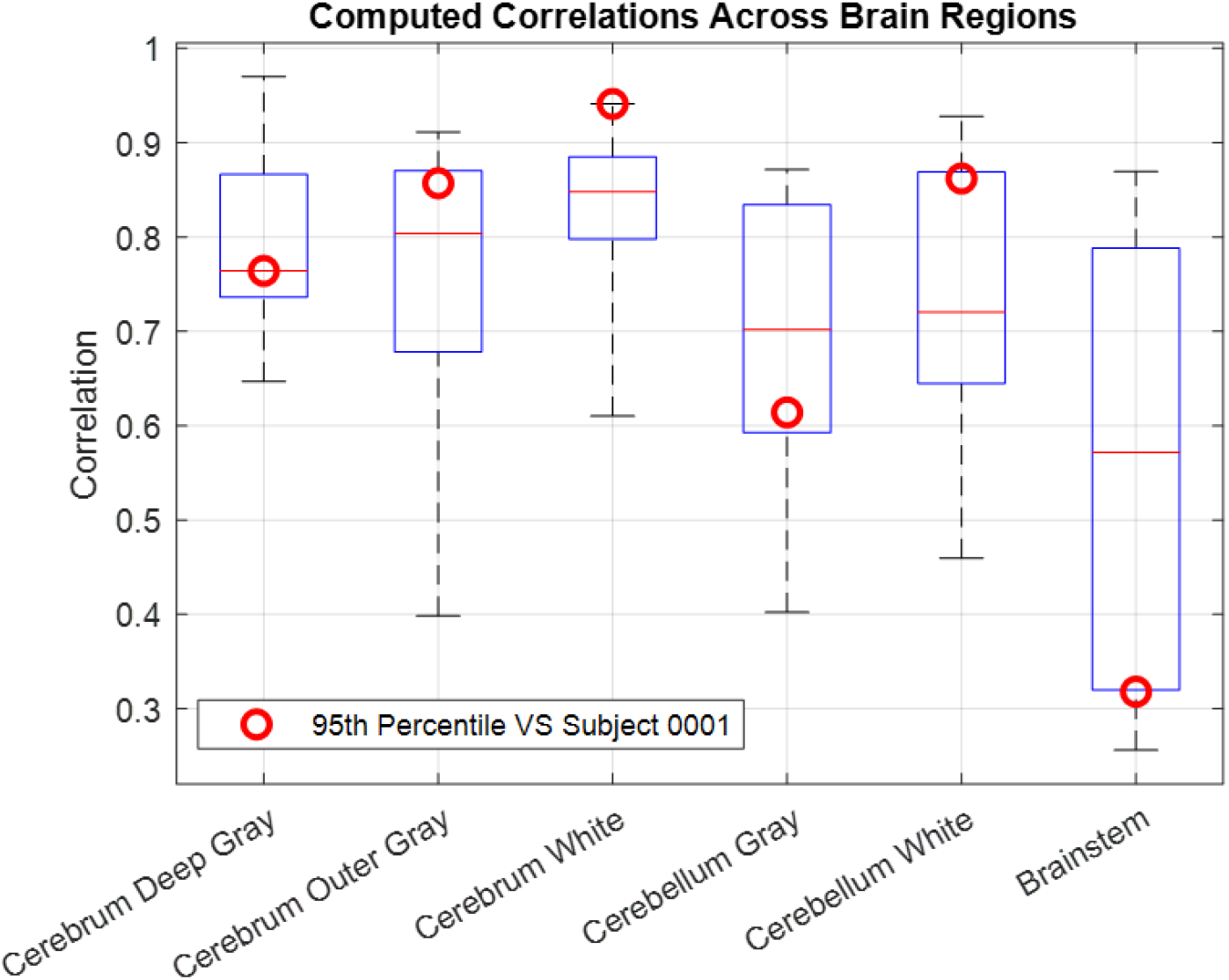
Correlations of the group average model. Red circles represent the Pearson correlations of the 95th percentile E_1_ between the NVHE group average model and subject 0001 across six brain segments. They share the same kinematic boundary condition. The blue boxes represent the 25th to 75th range of Pearson correlations of the 95th percentile E_1_ between the group average model and segments across all the mid male subjects. The whiskers cover the entire range of the Pearson correlations. The red lines are the medians of these Pearson correlations in different segments.

Furthermore, we can extend these correlations to the overall strain field of the group-average models by considering the following observations. Referring to Figure 13 in Appendix B, our first global observation is that E_1_ and E_3_ generally have similar magnitude but opposite signs (note the color bar), while E_2_ is close to zero throughout the times shown (27 *ms*, 45 *ms*, and 63 *ms*). Such similar magnitude and opposite signs are of course a consequence of the assumed incompressibility of the group-average model. Thus E_1_ is tensile while E_3_ is compressive, while E_2_ is generally negligible under this rotational type kinematic boundary condition. In this case, we can infer that knowing E_1_ allows us to infer all the three principal strains E_1_,E _2_, and E_3_, which is equivalent to knowing the entire strain tensor for this isotropic case. The same features can be observed by examining the subject-specific experimental data using imaging software ITK-SNAP. Therefore, knowing that our GAMS are representative in terms of E_1_ should ensure that the models are also representative in terms of E_3_ and E_2_.

A relatively common approach to modeling brain biomechanics is using linear material models, whether elastic or viscoelastic; these have the advantage of being computationally inexpensive and easy to calibrate. How well would such a model do in the case of these groups? The predictions of the linear viscoelastic (LVE) group-average model presented by Alshareef et al. [38] are shown as the purple lines in Figure. 4 and Figure 5 (the LVE simulations were computed using the commercial software LS-DYNA). The LVE simulations predict earlier peak times and larger strain magnitudes than the NVHE GAM and the experimental data across almost all brain segments. The LVE results also show a strong secondary peak near 70 ms which is not generally present in either the experimental or NVHE results. Further, note that the LVE results are very similar between the three segments shown in Figure 4, suggesting that the LVE model is not effective at differentiating the mechanical responses of these three brain segments. The pronounced secondary peaks also suggest that the LVE model does not well-capture the energy dissipation behavior. That said, the LVE model can still be useful as an upper-bound estimate of brain deformation for the particular low loading ranges considered in these experiments.

In Figure 7, we compare the spatial distribution of E_1_ in the brain of subject 0001 with the NVHE group-average model to demonstrate their common features in space. Again, we use subject 0001 for the comparison. The left three columns of Figure 7 show the E_1_ strain distributions for the six sections at *t* = 27 *ms*, 45 *ms*, and 63 *ms*, while the right 3 columns of Figure 7 are the corresponding (manually selected based on location, shape, and size) transverse planes from subject 0001 approximating the six group average transverse planes. Note that we exclude skull, ventricles, cerebrospinal fluid (CSF), and foramen of the group-average model for demonstration. Overall, the strains are seen to be propagating into the brain from the periphery in both the experimental data on this subject and the simulation results, while the local strain distributions are strongly dependent on the local geometry and material properties. We observe high strain regions (E_1_ >0.06) near the peripheries on slice 0 *mm*, 10 *mm*, and 15 *mm* at 45 *ms* for the both distributions. However, subject 0001 shows slight bilateral asymmetry in the strain distribution on these slices, which are not observed in the GAM simulations. This is perhaps attributable to experimental constraints: while group simulations assume idealized rotational boundary conditions, experimental neck rotations inevitably include minor translational motion, introducing center misalignment. The differences in the geometry are also immediately evident: compare the geometry of the slice at −30 *mm* between the model and the subject, for example. Even with similar geometry, such as on slice −27 *mm*, the group-average model shows a pronounced strain concentration in the sub-arachnoid space (modeled as LVE materials) at 45 *ms* and 63 *ms*, but was vaguely seen in the corresponding subject slices. With those caveats, we observe that the approximate magnitudes of the principal strains are similar for the model and the subject at the various times, but the spatial distributions of the strains are different in the detail.

**Figure 7:**
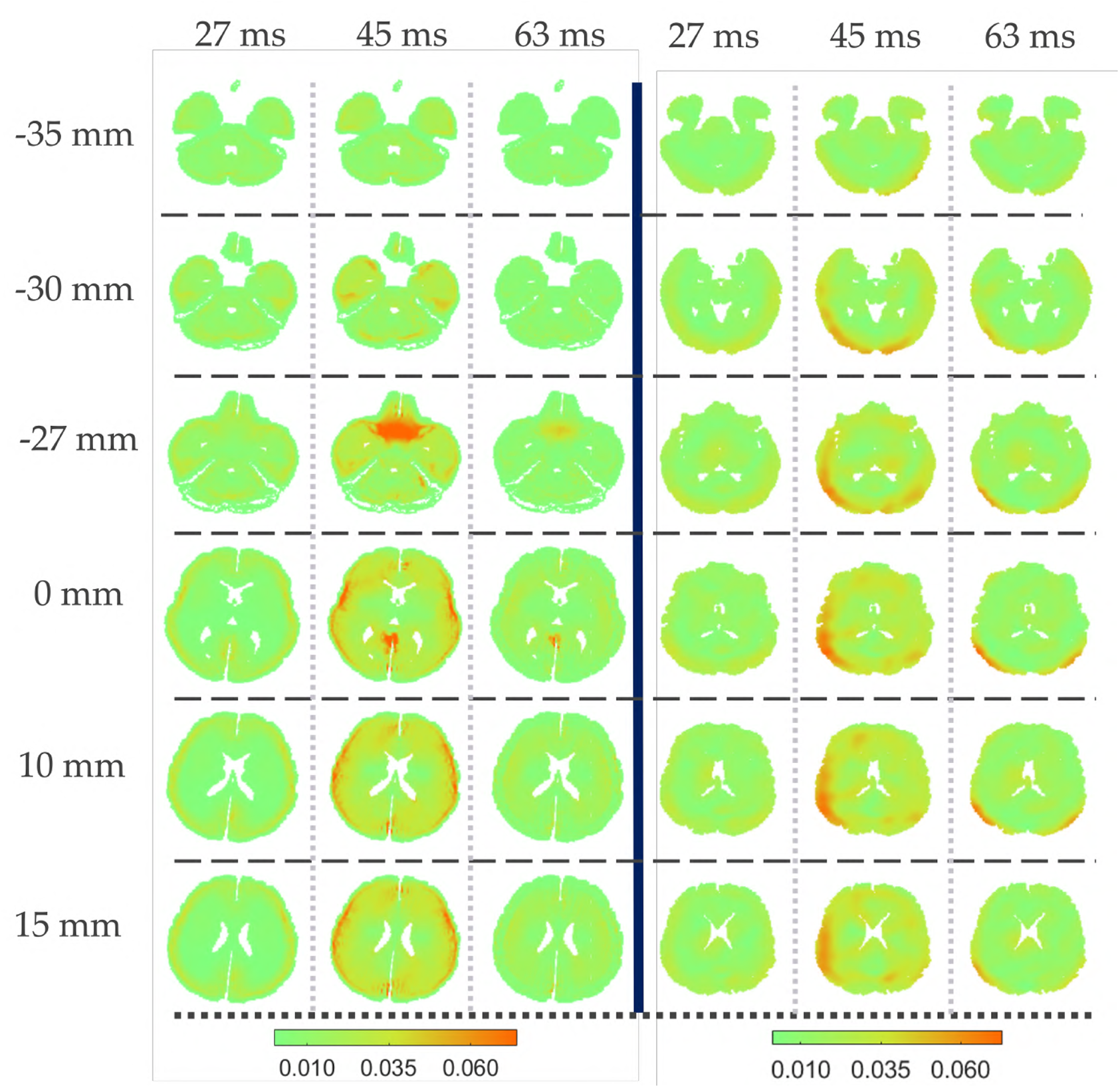
Spatial distribution of principal strain E_1_ of mid male group average simulation and a mid male E_1_ (left of the thick blue line) experimental data of subject 0001 (right of the thick blue line). Group average simulation is presented on six transverse slices with IS = −35 mm, −30 mm, −27 mm, 0 mm, 10 mm, and 15 mm at 27 ms, 45 ms, and 63 ms. Transverse planes of subject 0001 are manually selected based the its similarity of location, shape and size with the group average planes. The colorbars denote the magnitude of E_1_.

### 3.2 Differences in brain biomechanical responses of the six groups

Having established that the GAMs are representative of each group, we now seek to understand the differences in biomechanical response among the six groups. Note that our simulations with the six GAMs share the same kinematic boundary condition (from subject 0001), but the GAMs have different anatomical structures as described in section 2.1, and different material properties (Appendix A). To compare the simulation results from the six group-average models, we examine the entire brain volumes (thus excluding the skull, ventricles, CSF, and foramen), and create bins based on the magnitude of E_1_ within the entire brain at each time. We then plot the frequency of occurrence of each binned strain at a given time in a histogram plot for each of the six gender- and age-based GAMs. For each pair of groups (e.g. young female vs young male), we then plot comparative histograms, with the overlap regions identified (e.g., in Figure 8 examining gender-based differences). We perform these comparisons for a time of 45 *ms* (which generally corresponds to the time of the largest strain magnitudes in the strain histories).

**Figure 8:**
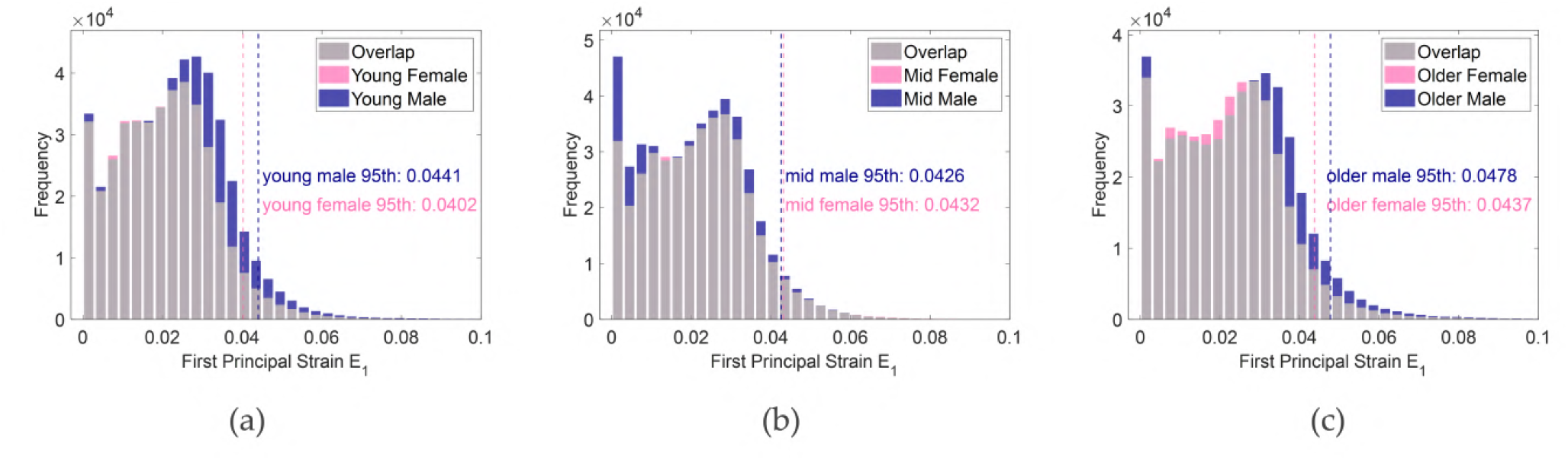
Histograms of all six groups at 45 ms. Each sub-figure fixes the age and compares histograms of different genders. 95th percentile of E_1_ are marked by vertical dashed lines.

These non-parametric histograms all show pronounced frequency peaks near zero strain, indicating that much of the brain is experiencing little deformation at 45 ms. This is because the applied kinematics boundary condition is a pulse with a loading duration of less than 50 ms, which is shorter than the characteristic time required for a shear wave to travel from the skull’s outer surface to the brain’s center [61]. We can also estimate the location of the wavefront: for example, from the static shear modulus G_*∞*_ of the brain white matter (corona radiata and corpus callosum) and the density of the brain, one can estimate the effective shear wavespeed is approximately 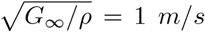. The distance from the periphery of the 0 *mm* transverse plane to the center of rotation is approximately 8 *cm* (obtained from the anatomical information), and so the shear wave would take 80 *ms* to reach the center. Thus, at 45 ms of the simulation, much of the brain volume still has not perceived the mechanical loading.

The histograms of Fig. 8 compare the results by gender for each age group: for instance, male (blue) compared to female (pink) in the mid-age group in Fig. 8 (b), and so forth. Overlaps between the two groups are plotted in gray. The 95th percentile E_1_ of each group is identified by drawing a vertical dashed line, with blue dash representing males and pink representing females. A global observation from this figure is that the male GAMs generally experience higher strains more frequently than the female GAMs, and this is most pronounced for the younger groups.

We quantify the differences through two-tailed p-tests of the 95th percentile E_1_ at 45 *ms*. As shown in Table 3, the two-tail hypothesis tests show that between young female/male, mid female/male, and older female/male, there are significant differences (*p* <0.001) in terms of the 95th E _1_ at 45 ms as the Z values are all much larger than 10 (p-test requires we choose a first-order statistic such as mean, 50th, and 95th percentile). Note that we choose the female as control group in each pair for *Z* − value computations. In Figure 8 (a), we find that the young male group has a greater 95th percentile E_1_ than young female, and more strains distributed between E_1_ of 0.02 and 0.06. Such a difference, as the computed *Z* − value shows, is highly unlikely (<0.001) to be caused by random errors. It appears that the differences are driven by the differences in the group average anatomical geometry and material properties. Overall, if one uses a maximum principal strain criterion to define onset of injury, Figure 8 suggests that males are more susceptible to injury than females, particularly in the younger age groups.

**Table 3:** Summary statistics by age and sex group.

| Group | 95th Percentile $E_1$ | Standard Deviation ($\sigma$) | No. Material Points $n$ | $\sqrt{n}$ | $Z$-value |
| --- | --- | --- | --- | --- | --- |
| Young Female | 0.04021 | 0.01217 | 403828 | 635.5 |  |
| Young Male | 0.04408 | 0.01325 | 475745 | 689.7 | 201 |
| Mid Female | 0.04318 | 0.01299 | 407680 | 638.5 |  |
| Mid Male | 0.04259 | 0.01326 | 454762 | 674.4 | -30 |
| Older Female | 0.04375 | 0.01374 | 392215 | 626.3 |  |
| Older Male | 0.04778 | 0.01504 | 431654 | 657.0 | 184 |

Note: $n$ is the number of material points excluding skull, ventricles, CSF, and foramen.

It is the case that from childhood to adulthood, the brains (and brain stiffnesses) of young males and females are still developing [62]. In Fig. 8 (b) we observe that the histograms of mid male and mid female groups show fewer differences than are observed for either the younger or the older groups. In Figure 8 (c), we find that the older male has a greater 95th percentile E_1_ than the older female, with more pronounced strain values between 0.03 and 0.06, which indicates that the older male brain GAM is also more susceptible to the same traumatic event than the older female brain GAM. This may relate to the observation that the male brain appears to shrink more than female brain during brain atrophy [63, 64, 65]. Appendix C compares the group-wise differences between ages of the same gender, showing consistency with these arguments.

These observations from Figure 8 raise a natural question: is it the difference in the group-wise anatomical geometry or the difference in the material properties that drives the differences in GAM behavior? To answer the question, we assemble a fictitious brain model called the “mix male,” using the mid male geometry with the older male material properties. We then compute the deformations of this “mix male” GAM under the same kinematic boundary condition as used for all of the GAM simulations. There are three reasons for considering this fictitious brain model. First, the mid-age groups (including both male and female, Table 1) have a greater number of MRI and MRE scans than the other groups, thus providing a more representative anatomical structure for our computations. Second, Figure 14(f) in Appendix C shows that the difference in strain histograms between the mid female and older female GAMs is much less than that between the mid male and older male GAMs (the material properties are also similar between mid and older females in our dataset, Table 6, at least considering the cerebrum segments which are up to 88% of the total brain volume [60]). Third, the material properties of cerebrum segments in the mid-male and older-male show approximately 10-20% difference. Now, comparing the results of simulations on the mid-male GAM and the mix-male GAM should tell us what the influence of the material properties is on these results. Note that Figure 9(b) shows that the mix-male GAM results are very similar to the older-male GAM results, showing that changing the anatomical structure alone produces very little difference in the results. However, Figure 9(a) shows clear differences in strain histograms frequency differences for all bins between the mid-male group and the mix-male group. Further, the 95th percentile E_1_ of the mix group 0.0508, is much closer to that of the older male group 0.0478. Putting these observations together, we conclude that the age-dependent change in material properties plays a greater role than the anatomical structure in driving the differences between the results of simulations using group average models.

**Figure 9:**
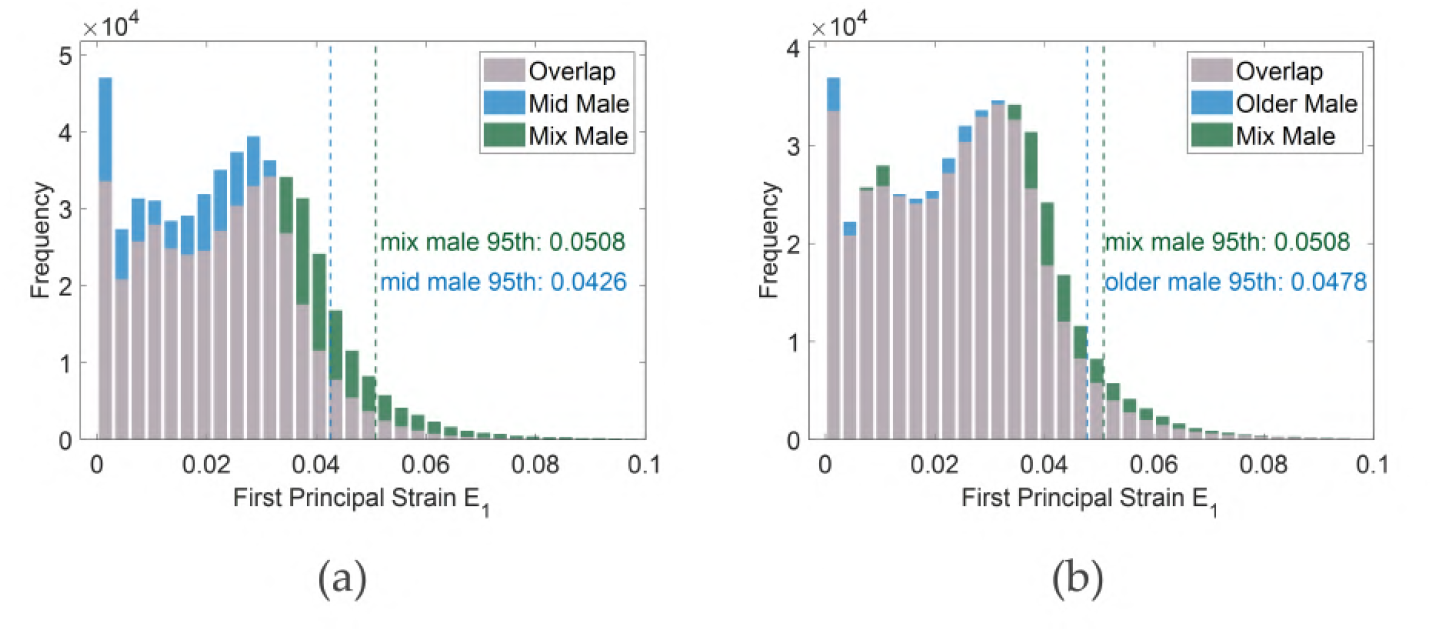
Histograms of (a) mixed male vs mid male group and (b) mixed male vs older male at 45 ms. 95th percentiles of E_1_ are marked by vertical dashed lines.

We compare the spatial differences in the strains between the six groups by selecting four transverse slices of the six groups at 45 *ms* for comparison. Consider the four slices of the mid-male group with IS coordinates of − 35 *mm*, − 27 *mm*, 0 *mm*, and 15 *mm* (Figure 3). We can find transverse planes of all of the six groups with similar IS coordinates, with some adjusted slightly according to the location, shape, and size of their own anatomical structure. We can now compare the strain fields for E_1_ of all the six groups at 45 ms across these four transverse planes, as shown in Figure 10. Anatomically, transverse slices of 15 mm show that the ventricles are significantly enlarged in the older groups, which is consistent with recent findings in [66, 67]. Such enlargement of ventricles may contribute to the overall decrease of the group-wise stiffness, perhaps causing the older groups to be more susceptible to a given traumatic insult. On the transverse slices of−27 *mm* of the six groups, we also notice that the size and magnitude of E_1_ in SAS are different across six groups. Note that the SAS (pointed by the black arrows in Figure 10) is modeled as viscoelastic, with the same parameters used across all of the six groups. We speculate that the differences in the SAS geometry, and the difference in geometry and material properties of nearby brain segments, cause the size and magnitude difference shown in the SAS. Note also that E_1_ in SAS and near falx are well over the 95th percentile strain shown in Figure 8. We will briefly discuss possible biomechanical explanations for this in the next section. Note that near the falx on the 0 *mm* plane of the young male GAM, a “checkerboarding” effect is evident, so that there are severe numerical errors in the region for this GAM at this time of 45 *ms*. More refined grids are needed to eliminate the checkerboarding.

**Figure 10:**
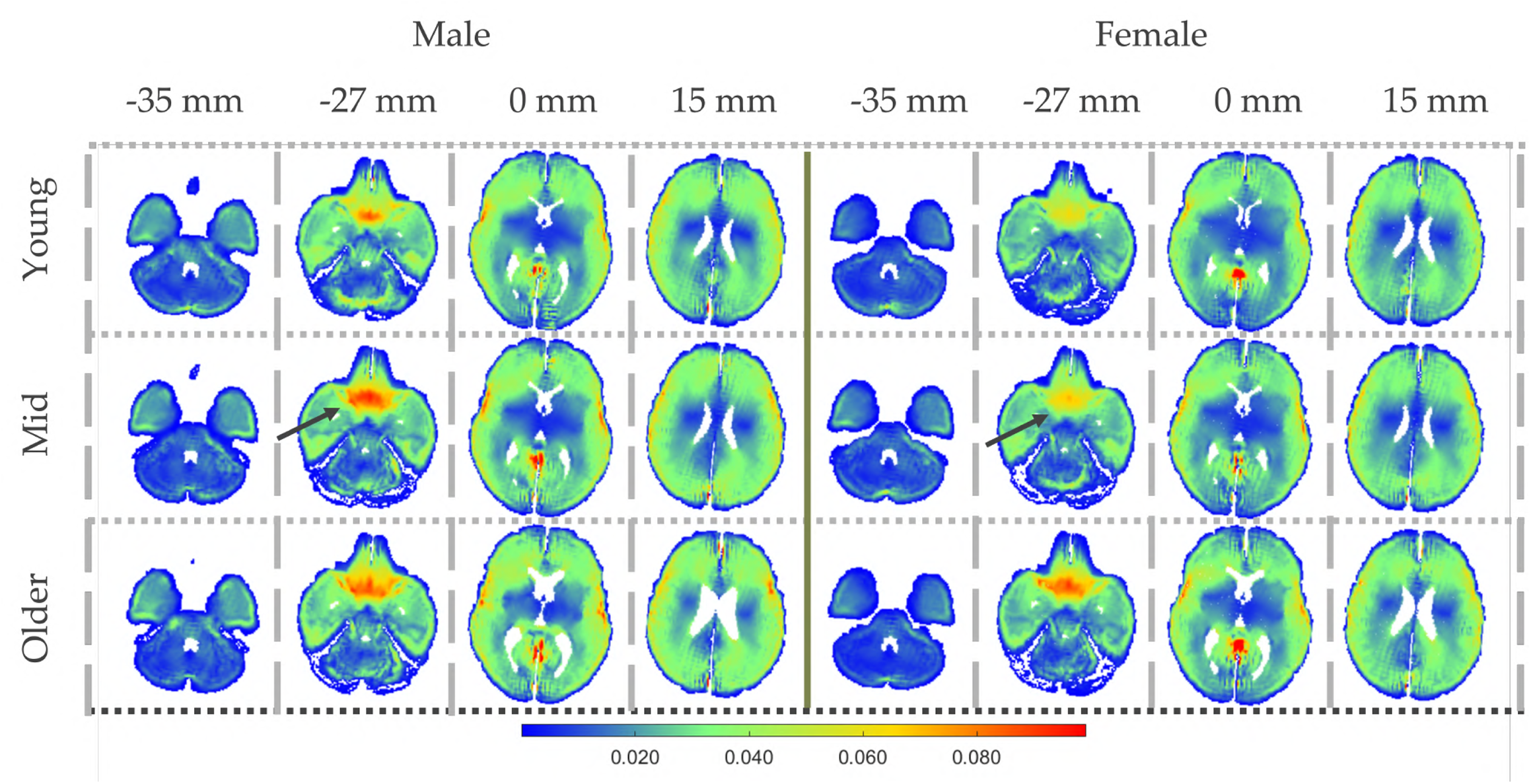
Spatial strain E_1_ distribution at 45 ms on four selected transverse planes of all six groups with kinematics boundary condition of subject 0001. The black arrows point to the SAS of mid male and mid female.

### 3.3 Estimation of Strain-Based Injury Criteria using Group-Average Models

We now use these established GAMs to examine the possible onset of injury under a potentially injurious loading condition, created by simply scaling the magnitude of the non-injurious kinematic boundary condition that we have applied previously to the GAMs. The new “injurious” kinematic boundary condition is defined as three times that applied to subject 0001 (i.e., three times the angular velocity and angular acceleration shown in Fig. 2). The scaling factor of three is selected primarily based on the experimental “conservative morphological injury threshold” of E_1_ = 0.14 used by Bain & Meaney [68], referring to the 95th percentile values of E_1_ observed in the cerebrum slice in Figure 4. This choice of scaling also provides similar order of magnitude with the experiments such as [69].

Several criteria have been proposed to identify potential injury locations under injurious loading, including strain thresholds, strain rate thresholds, and combinations of these metrics [70, 69, 71]. In this study, we consider both a strain-based injury criterion as noted above, as well as a strain-rate–based criterion with a threshold of 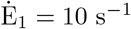 [72, 73, 74], which is considered as a finite (rather than *infinitesimal* rate in works such as [75, 76]. Note that because these injury thresholds take finite rather than *infinitesimal* values, the accurate evaluation of injury necessitates the use of finite deformation (nonlinear) constitutive models such as the nonlinear visco-hyperelastic model that we use here.

We apply this injurious kinematic boundary condition to all six GAMs, and estimate the locations and magnitudes of strain-based injury within these brains using the two thresholds identified above. The results are presented in Figure 11 by highlighting the injured regions across the six groups at the time of the peak strain, 45 *ms*. The blue highlighted regions are those with E _1_ >0.14; the orange regions are those with 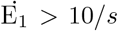 and the regions having both thresholds satisfied are highlighted in purple. We use these highlighted regions to identify the specific regions of the brain that are vulnerable to injury under this specific loading for each of the six age and gender-stratified groups.

**Figure 11:**
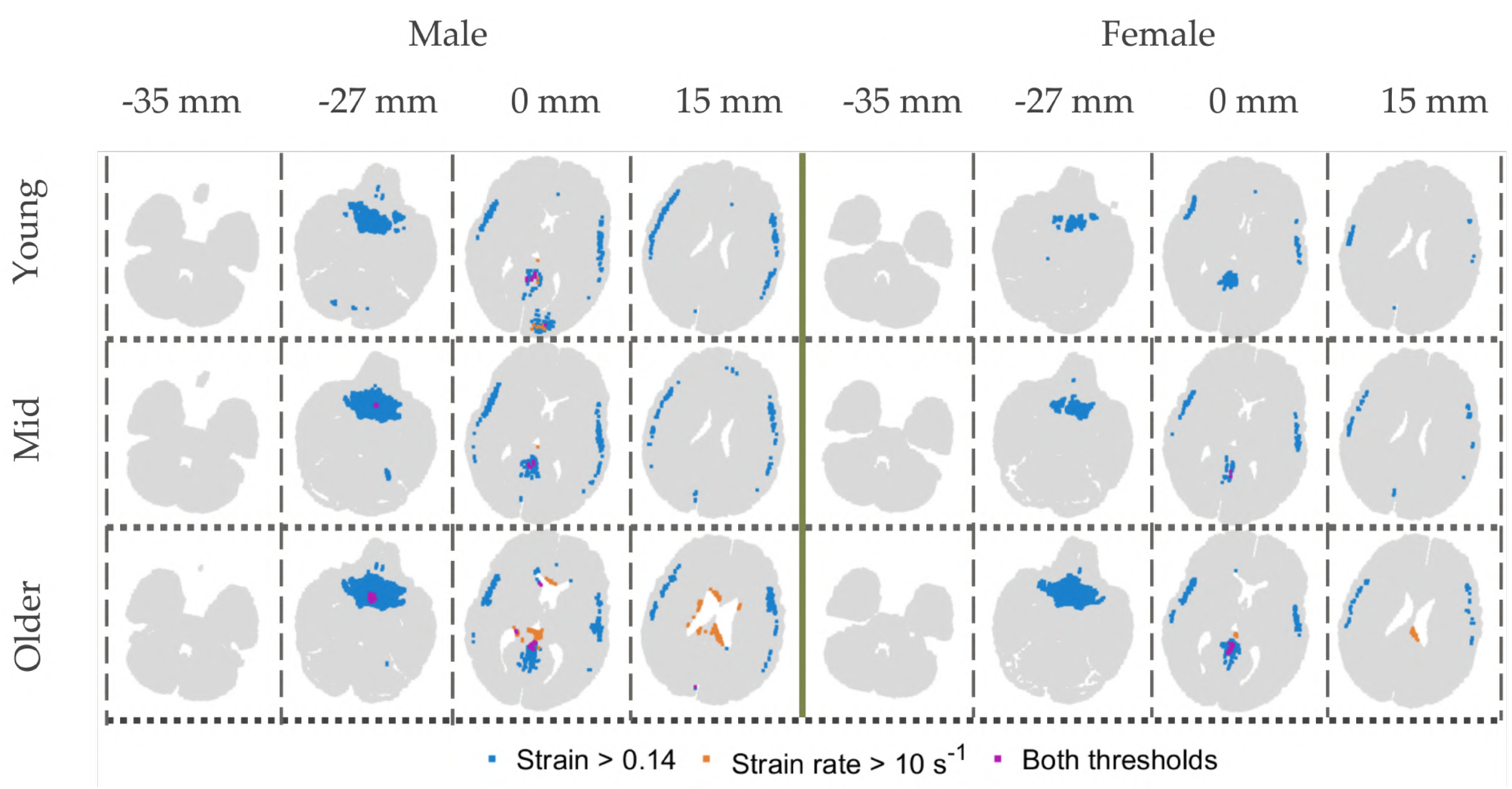
Vulnerable regions under injurious loading of all six groups at 45 ms with strain and strain rate threshold of 0.14 and 10 s^−1^, respectively.

The critical-strain-based vulnerable regions of all six groups are located in the cerebrum outer regions and the brain tissues near the falx. The critical-strain-rate-based vulnerable regions are observed in all male groups, but only in the older female under this loading condition. The critical-strain-rate-based regions are mainly located near the interfaces of brain segments such as corpus callosum near the falx and ventricle. The overlaps between the two types of injured regions are mainly located near the falx (i.e., if both criteria need to be satisfied simultaneously). Note that the peak magnitude of strain rate does not synchronize with the peak magnitude of strain in our results; the maximum strain rate occurs before 45 *ms* in general. Readers that are interested in the full field strain distributions of all the six groups on the same four transverse slices at 45 *ms* are directed to Figure 15 of Appendix D (Note that the colorbar has a different scale there compared to Figure 10). All of these results will be available in a publicly available dataset.

We note that the fidelity of our GAM with respect to the behavior of the SAS is low, since in modeling SAS, what material constitutive relations and the corresponding parameters to use are still an open question [46]. To the best of our knowledge, there is no agreement on what strain values can be considered as thresholds for the SAS. With those caveats, injury mechanisms like subarachnoid hemorrhage (SAH) following TBI [77] have been identified in the SAS, and this should perhaps be considered in future work using such GAMs. High strain regions in SAS may be due also to the relative low modulus assigned (refer to in table 4 and Appendix A).

The high strain values in the cerebrum outer regions on transverse slices 0 *mm* and 15 *mm* are the direct consequence of strain concentration due to material modulus mismatch at the interface of the cerebrum outer gray and cerebrum white regions, and the fact that the cerebrum outer gray matter is the first layer of the brain tissue that is subjected to the injurious loading. These regions also correspond to locations that are identified for commonly diagnosed diffuse axonal injury (DAI), such as the corticomedullary injunction [78]. In the older male and older female groups, the “injured” (rate-threshold) regions on the 15 *mm* transverse slice are part of the corpus callosum near the ventricles, which is another common location in DAI diagnoses [78].

The “injured” blue (strain-threshold) region near the falx (note that falx is excluded in Fig. 11) is likely also affected by material interfaces. When the shear stress wave in the brain encounters a much stiffer material at the falx (see Table 2), we expect the wave reflections to result in the accumulation of strain in the adjacent cerebrum tissues. Note that in our group-average computational models, we assume perfectly bonded interfaces [53]. Large strains near the falx cerebri may also cause falx displacement, leading to subfalcine herniation [79], and inter-hemispheric subdural hematoma [80].

## 4 Key limitations of the work

Our research has several significant limitations. First, our young male and young female group has ages in the range of 18-21, resulting in two potential issues. Human brains in this age range may still be under development, during which (e.g.) the brain stiffness may undergo significant changes [62]. Further, such an age grouping assigns people of age 22 to 30 to the mid age groups, which may be unreasonable for the lower ages. Future studies could perhaps consider age grouping criteria based on the development of the human brain. Second, our models have limited fidelity and resolution with respect to the anatomical structure and material properties. With respect to anatomical structure, we need more refined segmentation in brain membrane structures such as tentorium and falx. Such refinement would allow further differentiation of material properties in the regions of interest, such as anterior and posterior falx. With respect to material properties, our current GAMs do not incorporate fiber orientations and fractional anisotropy, making our models unable to predict axonal strains. The next generation group-average models should incorporate these two factors as they are essential for predicting diffuse axonal brain injuries. Third, the current models use a grid resolution of 96 × 96 × 72. Under this resolution, we can identify high strain regions near material interfaces, but we cannot resolve fine-scale strain gradients at the material interfaces. However, such interfaces are of great concern in clinical practice. As always, computational models such as these would benefit from greater refinement and additional materials data, and there are tradeoffs involved in exploration of the landscape versus high fidelity high resolution simulations of specific conditions.

## 5 Concluding remarks

This study demonstrates that age- and gender-stratified group-average models can effectively represent biomechanical responses of the individuals within the groups, and can reveal meaningful demographic differences that may influence susceptibility to traumatic brain injury. We have demonstrated (in detail for one group) that our group average computational models can represent, to reasonable degree, the brain biomechanics associated with individual subjects in the group. We compute the Pearson correlations between the group-average models and the subjects in the group across different brain segments, and find strong correlation (Pearson correlation> 0.7) in most brain segments, and moderate correlation in some segments. We also find that there are significant differences (*P* <0.001) across all the six groups average models (at least in terms of the maximum principal strain E_1_), indicating that there is value in such age-and-gender-stratified grouping of biomechanical data. To determine the relative importance of the group-average geometry and material properties, we conducted a numerical experiment by assembling the mid male group geometry and older male material properties, and find that the material properties are more dominant in determining the strain differences. Finally, we apply an injurious loading to our models, and demonstrate their efficacy in revealing vulnerable regions of the brain with respect to this kinematic boundary condition. We hope to see increased utilization of these group-average models in both research and clinical applications.

## 6 Acknowledgements

This work was partially supported by NIH grant U01NS112120 (P.V. Bayly, K.T. Ramesh, J. Prince, D. Pham, and C. Johnson). We would like to thank Dr. Kshitiz Upadhyay and Dr. Yuan-Chiao Lu for discussions during the research.

## Appendix A: O-USS material parameters of Other Group Average Models

**Table 4:** Young male group average O-USS material parameters.

| O-USS material parameters |  |  |  |  |  |  |  |
| --- | --- | --- | --- | --- | --- | --- | --- |
| Brain Substructure | $G_\infty$ (Pa) | $\alpha$ | $k_{11}$ (Pa·s) | $k_{21}$ (Pa·s<sup>2-2c<sub>21</sub></sup>) | $c_{21}$ | $\kappa$ (GPa) | $\rho$ (kg/m<sup>3</sup>) |
| Brain Stem | 676.4 | 2.61 | 1.6 | 263.7 | 0.588 | 1.46 | 1040 |
| Cerebellar Gray | 858.6 | 2.71 | 7.7 | 335.2 | 0.62 |  |  |
| Cerebellar White | 1092.8 | 2.71 | 5.5 | 490 | 0.605 |  |  |
| Corona Radiata | 1147.2 | -3.47 | 4.4 | 595.8 | 0.605 |  |  |
| Corpus Callosum | 1315.4 | -2.32 | 5.2 | 562.9 | 0.58 |  |  |
| Cortical Gray | 1011.5 | -3.76 | 6.3 | 436.9 | 0.622 |  |  |
| Deep Gray | 1425.7 | 4.92 | 7.3 | 694.7 | 0.601 |  |  |

**Table 5:** Young female group average O-USS material parameters.

| O-USS material parameters |  |  |  |  |  |  |  |
| --- | --- | --- | --- | --- | --- | --- | --- |
| Brain Substructure | $G_\infty$ (Pa) | $\alpha$ | $k_{11}$ (Pa·s) | $k_{21}$ (Pa·s<sup>2-2c<sub>21</sub></sup>) | $c_{21}$ | $\kappa$ (GPa) | $\rho$ (kg/m<sup>3</sup>) |
| Brain Stem | 1146.4 | 2.61 | 2.3 | 385.8 | 0.569 | 1.46 | 1040 |
| Cerebellar Gray | 644.0 | 2.71 | 11.3 | 282.4 | 0.658 |  |  |
| Cerebellar White | 836.5 | 2.71 | 10.1 | 418.7 | 0.627 |  |  |
| Corona Radiata | 1165.2 | -3.47 | 6.5 | 612.7 | 0.621 |  |  |
| Corpus Callosum | 1394.4 | -2.32 | 5.1 | 580.5 | 0.576 |  |  |
| Cortical Gray | 964.4 | -3.76 | 10.0 | 440.5 | 0.640 |  |  |
| Deep Gray | 1534.8 | 4.92 | 6.9 | 678.3 | 0.588 |  |  |

**Table 6:** Mid female group average O-USS material parameters.

| O-USS material parameters |  |  |  |  |  |  |  |
| --- | --- | --- | --- | --- | --- | --- | --- |
| Brain Substructure | $G_\infty$ (Pa) | $\alpha$ | $k_{11}$ (Pa·s) | $k_{21}$ (Pa·s<sup>2-2c<sub>21</sub></sup>) | $c_{21}$ | $\kappa$ (GPa) | $\rho$ (kg/m<sup>3</sup>) |
| Brain Stem | 954.8 | 2.61 | 3.5 | 386.5 | 0.592 | 1.46 | 1040 |
| Cerebellar Gray | 671.5 | 2.71 | 9.6 | 313.7 | 0.640 |  |  |
| Cerebellar White | 978.0 | 2.71 | 9.7 | 490.9 | 0.621 |  |  |
| Corona Radiata | 1167.0 | -3.47 | 5.9 | 615.5 | 0.618 |  |  |
| Corpus Callosum | 1285.3 | -2.32 | 5.6 | 487.6 | 0.578 |  |  |
| Cortical Gray | 898.1 | -3.76 | 10.2 | 421.7 | 0.643 |  |  |
| Deep Gray | 1424.9 | 4.92 | 7.0 | 650.3 | 0.590 |  |  |

**Table 7:** Older male group average O-USS material parameters.

| O-USS material parameters |  |  |  |  |  |  |  |
| --- | --- | --- | --- | --- | --- | --- | --- |
| Brain Substructure | $G_\infty$ (Pa) | $\alpha$ | $k_{11}$ (Pa·s) | $k_{21}$ (Pa·s<sup>2-2c<sub>21</sub></sup>) | $c_{21}$ | $\kappa$ (GPa) | $\rho$ (kg/m<sup>3</sup>) |
| Brain Stem | 782.3 | 2.61 | 1.3 | 289.8 | 0.569 | 1.46 | 1040 |
| Cerebellar Gray | 852.1 | 2.71 | 6.0 | 344.6 | 0.607 |  |  |
| Cerebellar White | 1224.5 | 2.71 | 3.7 | 471.9 | 0.571 |  |  |
| Corona Radiata | 1036.7 | -3.47 | 4.8 | 535.5 | 0.611 |  |  |
| Corpus Callosum | 1336.4 | -2.32 | 3.4 | 490.6 | 0.564 |  |  |
| Cortical Gray | 848.9 | -3.76 | 8.2 | 371.2 | 0.640 |  |  |
| Deep Gray | 1412.0 | 4.92 | 4.3 | 320.5 | 0.573 |  |  |

**Table 8:** Older female group average O-USS material parameters.

| **O-USS material parameters** |  |  |  |  |  |  |  |
| --- | --- | --- | --- | --- | --- | --- | --- |
| Brain Substructure | $G_\infty$ (Pa) | $\alpha$ | $k_{11}$ (Pa·s) | $k_{21}$ (Pa·s<sup>2-2c<sub>21</sub></sup>) | $c_{21}$ | $\kappa$ (GPa) | $\rho$ (kg/m<sup>3</sup>) |
| Brain Stem | 717.3 | 2.61 | 1.9 | 284.9 | 0.587 | 1.46 | 1040 |
| Cerebellar Gray | 698.0 | 2.71 | 10.7 | 597.3 | 0.641 |  |  |
| Cerebellar White | 1014.7 | 2.71 | 10.6 | 447.0 | 0.622 |  |  |
| Corona Radiata | 1012.0 | -3.47 | 5.9 | 522.8 | 0.621 |  |  |
| Corpus Callosum | 1319.8 | -2.32 | 2.9 | 427.5 | 0.552 |  |  |
| Cortical Gray | 828.6 | -3.76 | 8.5 | 369.0 | 0.638 |  |  |
| Deep Gray | 1284.9 | 4.92 | 6.4 | 592.8 | 0.595 |  |  |

## Appendix B: Supplemental materials for group-average representativeness

**Figure 12:**
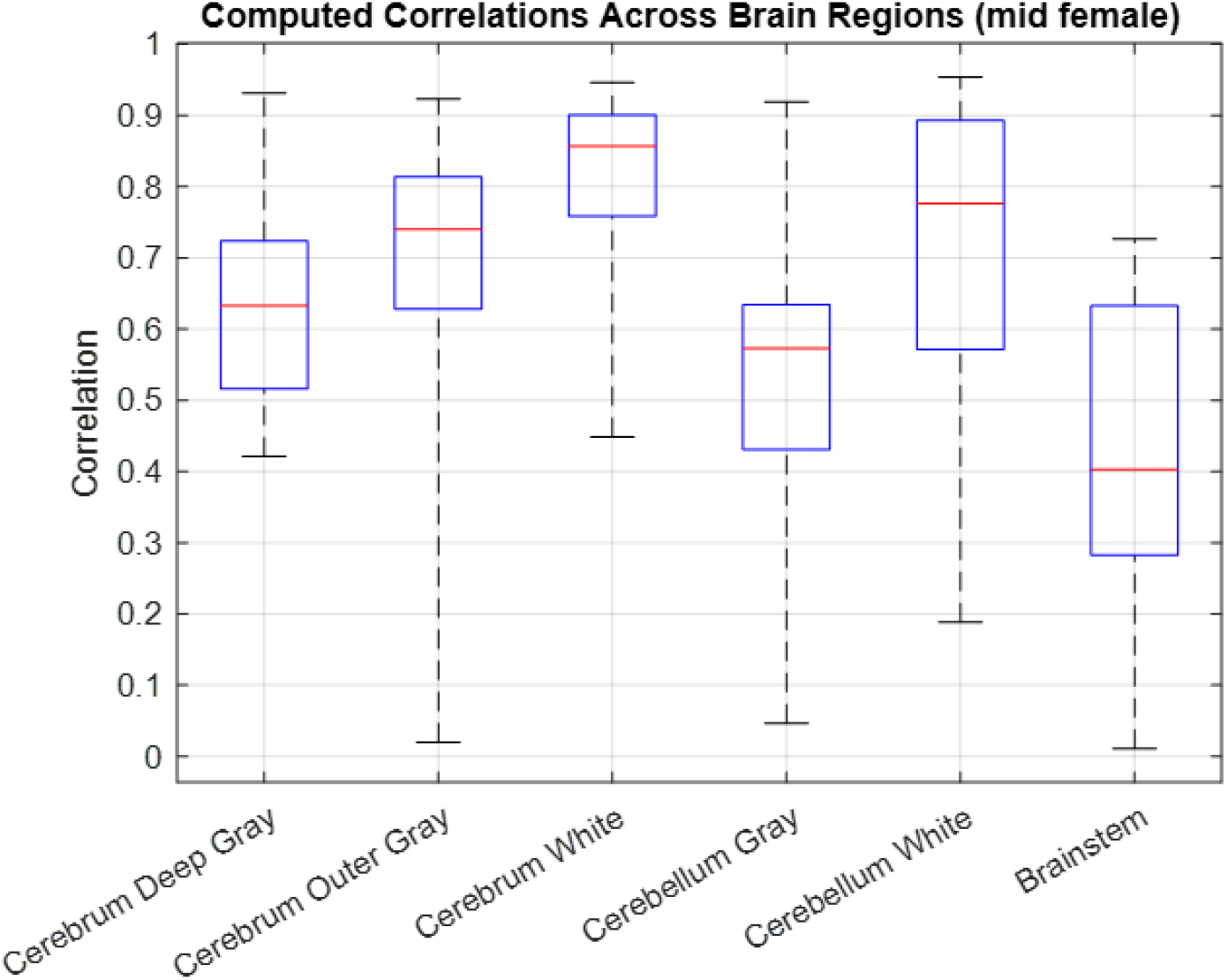
Correlations of the group average model of mid female. The blue boxes represent the 25th to 75th range of Pearson correlations of the 95th percentile E_1_ between the group average model and segments across all the mid male subjects. The whiskers cover the entire range of the Pearson correlations. The red lines are the medians of these Pearson correlations in different segments.

**Figure 13:**
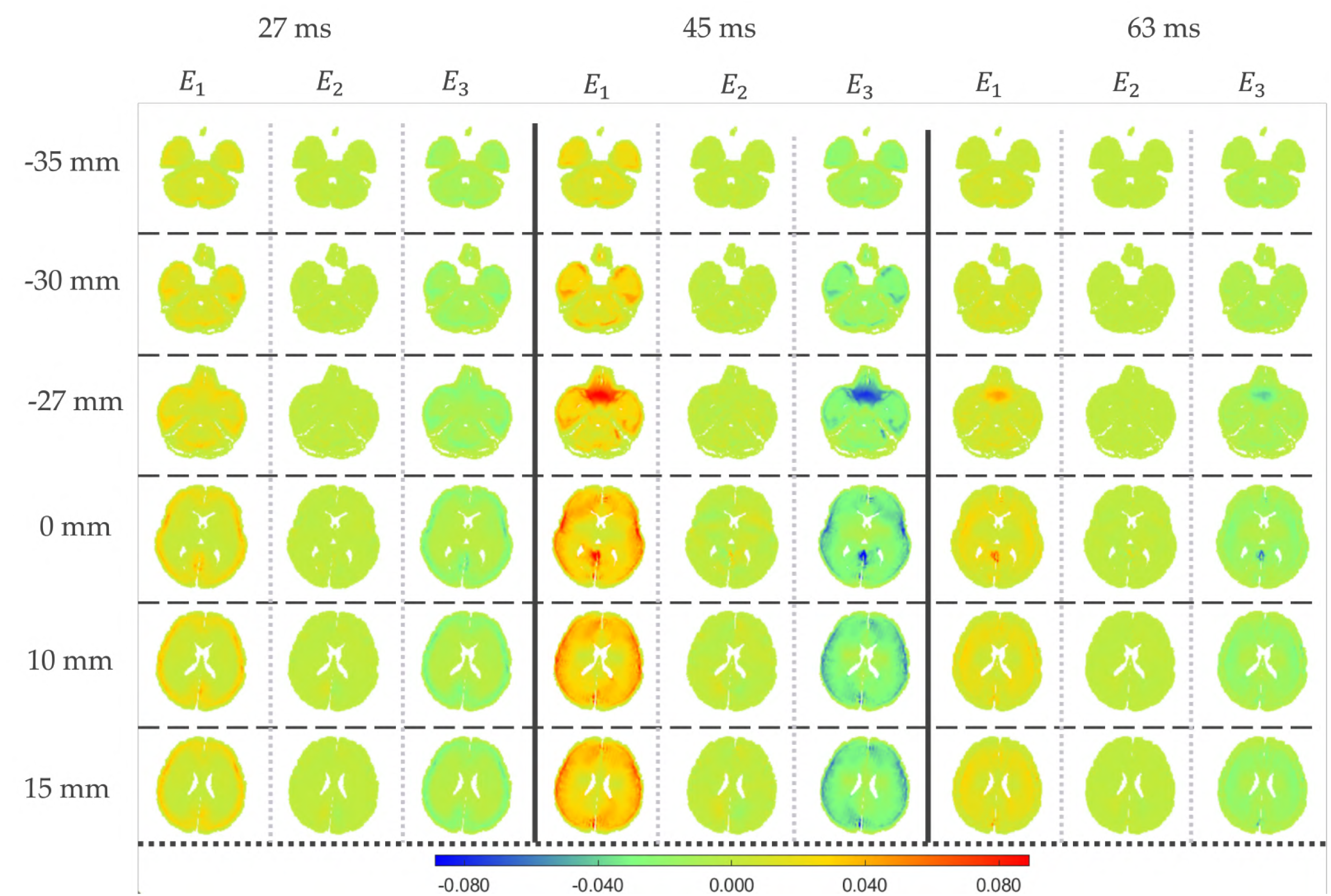
Spatial distribution of the group average model of E_1_,E _2_, and E_3_ on six transverse slices with IS = −35 *mm*, −30 *mm*, −27 *mm*, 0 *mm*, 10 *mm*, and 15 *mm* at 27 *ms*, 45 *ms*, and 63 *ms*. Note that the skull, ventricles, cerebrospinal fluid (CSF), and foramen are omitted for the purposes of this visualization.

## Appendix C: Supplemental materials for group-wise differences

**Figure 14:**
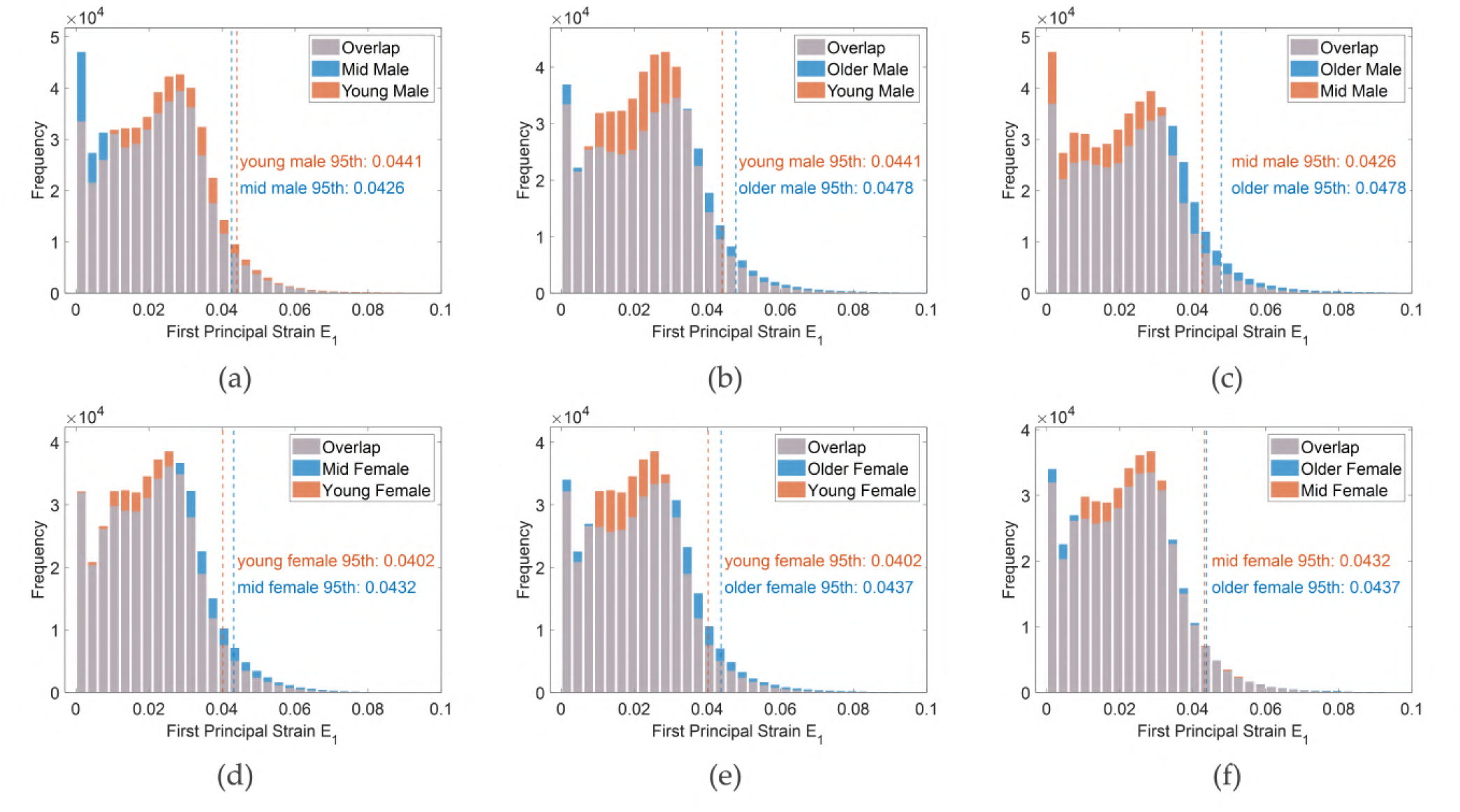
Histograms that fix gender and compare different age groups. 95th percentile of E_1_ are marked by vertical dashed lines.

## Appendix D: Strain distribution of six groups under injurious loading

**Figure 15:**
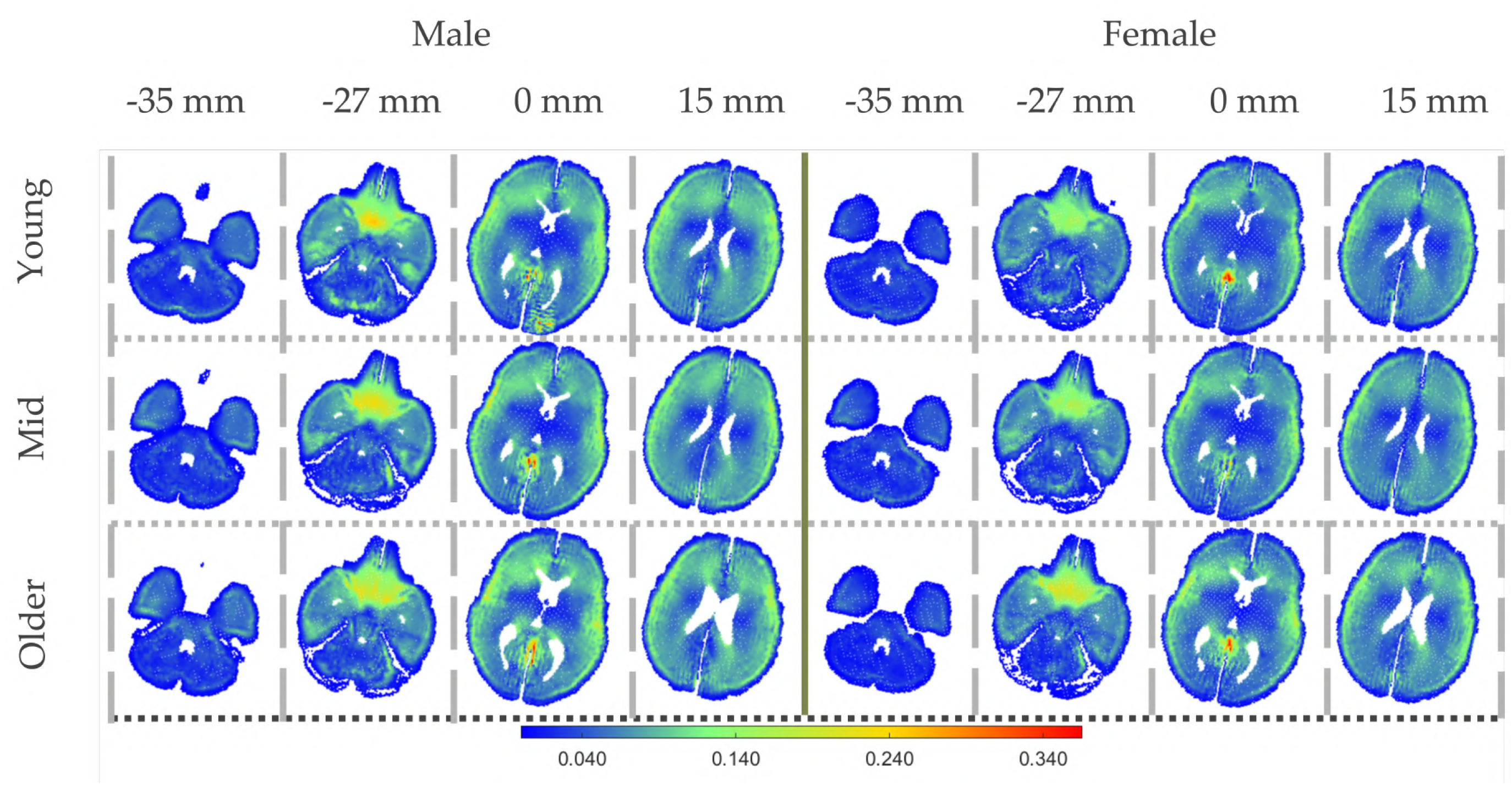
Spatial strain E_1_ distribution at 45 ms on six selected transverse planes of all six groups with 3 times kinematic boundary condition (injurious loading). Note that the strain beyond 0.35 may be caused from MPM numerical errors when the Lagrangian material points pass the cell grids [81, 82] and the material points are located at the material interface [83, 84].

